# NNT Regulates Mitochondrial Metabolism in NSCLC Through Maintenance of Fe-S Protein Function

**DOI:** 10.1101/761577

**Authors:** Nathan P. Ward, Yun Pyo Kang, Aimee Falzone, Terry A. Boyle, Gina M. DeNicola

**Affiliations:** Department of Cancer Physiology, Moffitt Cancer Center, Tampa, FL; Department of Molecular Pathology, Moffitt Cancer Center, Tampa, FL

**Author notes:** <u>**Summary**</u> Human lung tumors engage in substantial mitochondrial metabolism. Here, Ward et al. demonstrate that NNT supports oxidative metabolism in NSCLC through its mitigation of Fe-S cluster oxidation.

## Abstract

Human lung tumors exhibit robust and complex mitochondrial metabolism, likely precipitated by the highly oxygenated nature of pulmonary tissue. As ROS generation is a byproduct of this metabolism, reducing power in the form of nicotinamide adenine dinucleotide phosphate (NADPH) is required to mitigate oxidative stress in response to this heightened mitochondrial activity. Nicotinamide nucleotide transhydrogenase (NNT) is known to sustain mitochondrial antioxidant capacity through the generation of NADPH, however its function in non-small cell lung cancer (NSCLC) has not been established. We found that NNT expression significantly enhances tumor formation and aggressiveness in mouse models of lung tumor initiation and progression. We further show that NNT loss elicits mitochondrial dysfunction independent of substantial increases in oxidative stress, but rather marked by the diminished activities of proteins dependent on resident iron-sulfur clusters. These defects were associated with both NADPH availability and ROS accumulation, suggesting that NNT serves a specific role in mitigating the oxidation of these critical protein cofactors.

## Introduction

Metabolic rewiring facilitates the diversion of intermediate metabolites into pathways that supply the macromolecular determinants of the unbridled growth associated with human tumors. It is now appreciated that in addition to hallmark Warburg metabolism, many tumor species require substantial mitochondrial metabolism to thrive (Porporato et al., 2018). Indeed, non-small cell lung cancer (NSCLC) exhibits a simultaneous engagement of both glycolytic and oxidative metabolism (Fan et al., 2009). This increase in mitochondrial oxidation is supported by enhanced expression of pyruvate carboxylase, which provides auxiliary entry of glucose carbon into the TCA cycle in addition to that via pyruvate dehydrogenase (PDH) (Sellers et al., 2015, Davidson et al., 2016). These activities support glucose oxidation in human lung tumors that exceeds that of normal adjacent lung (Hensley et al., 2016). Moreover, human lung tumors exhibit remarkable plasticity in oxidative fuel usage that is correlated with oxygen tension, indicating robust mitochondrial function (Hensley et al., 2016, Faubert et al., 2017).

Mitochondrial redox metabolism harnesses the potential energy stored in the reducing equivalents nicotinamide adenine dinucleotide (NADH) and flavin adenine dinucleotide (FADH_2_), which are generated from the catabolism of various carbon substrates (i.e. glucose, amino acids, fatty acids). These are then oxidized by multiprotein complexes of the electron transport chain (ETC), which couples the transfer of electrons to molecular oxygen with the generation of a proton gradient. This proton gradient is exploited by ATP synthase to generate useable energy for the cell in the form of ATP. Critical to ETC functionality is the maintenance of mitochondrial reducing power in the form of nicotinamide adenine dinucleotide phosphate (NADPH). NADPH is required for the detoxification of harmful ROS that are natural byproducts of oxidative metabolism (Navarro et al., 2017). ROS detoxification prevents the oxidation of proteins critical to metabolism, including the four respiratory chain protein complexes (I-IV) of the ETC, as well as other macromolecules such as the constituent lipid species of the inner mitochondrial membrane (IMM). Additionally, ROS detoxification protects against the oxidation of iron-sulfur (Fe-S) clusters, redox sensitive cofactors assembled within the mitochondria and incorporated into recipient Fe-S proteins that facilitate diverse functions such as DNA replication and repair, protein translation, and metabolism (Flint et al., 1993, Alhebshi et al., 2012, Rouault, 2015). Mitochondrial metabolism is particularly dependent on these clusters as fatty acid catabolism, respiration, and cofactor synthesis (e.g lipoic acid, heme) all rely on Fe-S proteins (Lill and Muhlenhoff, 2008).

There are several sources of mitochondrial NADPH, including serine-dependent one-carbon metabolism (Ducker et al., 2016), isocitrate dehydrogenase (Jiang et al., 2016), malic enzyme (Ren et al., 2014) and the nicotinamide nucleotide transhydrogenase (NNT). NNT is an integral membrane protein associated with the IMM that harnesses the proton gradient across the membrane to couple the oxidation of NADH to the reduction of NADP^+^, yielding NADPH (Rydstrom, 2006, Kampjut and Sazanov, 2019). In bacteria, transhydrogenase activity accounts for up to 45% of total NADPH, which has led to the assertion that NNT should be an equal or greater contributor to the mitochondrial pool in mammalian species (Sauer et al., 2004, Rydstrom, 2006). Indeed, in the liver of mice, nearly 50% of the mitochondrial NADPH pool is sensitive to IMM uncoupling, indicative of a significant role for NNT (Klingenberg and Slenczka, 1959). Through subsequent studies NNT has been established as a significant contributor to the mitochondrial NADPH pool (Fisher-Wellman et al., 2015, Ronchi et al., 2016, Francisco et al., 2018). The importance of NNT has also been evaluated in malignancy, where NNT is shown to contribute to the maintenance of redox homeostasis in several cancers (Gameiro et al., 2013, Ho et al., 2017, Chortis et al., 2018, Li et al., 2018).

While the maintenance of mitochondrial NADPH has been shown to be critical to lung tumor growth (Ren et al., 2014, Jiang et al., 2016), the contribution of NNT to lung tumorigenesis has not been evaluated. Herein, we provide the first evidence that NNT is a significant contributor to the mitochondrial NADPH pool in NSCLC and important for lung tumorigenesis. We show that loss of NNT activity disrupts mitochondrial metabolism in part through diminished Fe-S protein function. Our data indicate a more nuanced role for NNT in NSCLC redox homeostasis through the prevention of Fe-S cluster oxidation rather than protecting against global mitochondrial oxidation.

## Results

### NNT Supports Lung Tumorigenesis

Many common conditional lung tumor mouse models were generated from breeding strategies that employed C57BL6/J mice (Jackson et al., 2001, Jackson et al., 2005, Meuwissen and Berns, 2005). Interestingly, these mice carry a homozygous in-frame deletion of exons 7-11 as well as a missense mutation in the mitochondrial leader sequence of the *NNT* gene that result in the expression of a non-functional protein (Toye et al., 2005). Therefore, to assess the contribution of NNT to lung tumorigenesis, we exploited this natural knockout allele.

First, we used a lung tumor model driven by mutant Kras (*LSL-Kras*^*G12D*/+^) to examine the influence of NNT expression on lung tumor initiation. Infection of *LSL-Kras*^*G12D*/+^ mice with adenovirus encoding Cre recombinase induces the expression of Kras^G12D^ in the lung epithelium and the formation of lung adenomas that rarely progress to higher grade tumors (Jackson et al., 2001). We found that expression of NNT in this model resulted in significantly greater tumor burden 3-months following Cre recombinase induction (Figs. 1 A and 1 B). Next, we used a lung tumor progression model that is driven by concomitant expression of mutant Kras and p53 deletion (*LSL-Kras*^*G12D*/+^; *Trp53*^*flox/flox*^, aka KP). NNT expression did not alter the survival of KP mice following Cre-induction (Fig. 1 C). Interestingly, p53 deletion abrogated the effects of NNT expression on lung tumor formation in this model, with substantial tumor burden present at the experimental endpoint regardless of NNT status (Fig. 1 D). Quantification of the fraction of burdened lung demonstrated no difference across genotypes (Figs. 1 E and 1 F).

**Figure 1.**
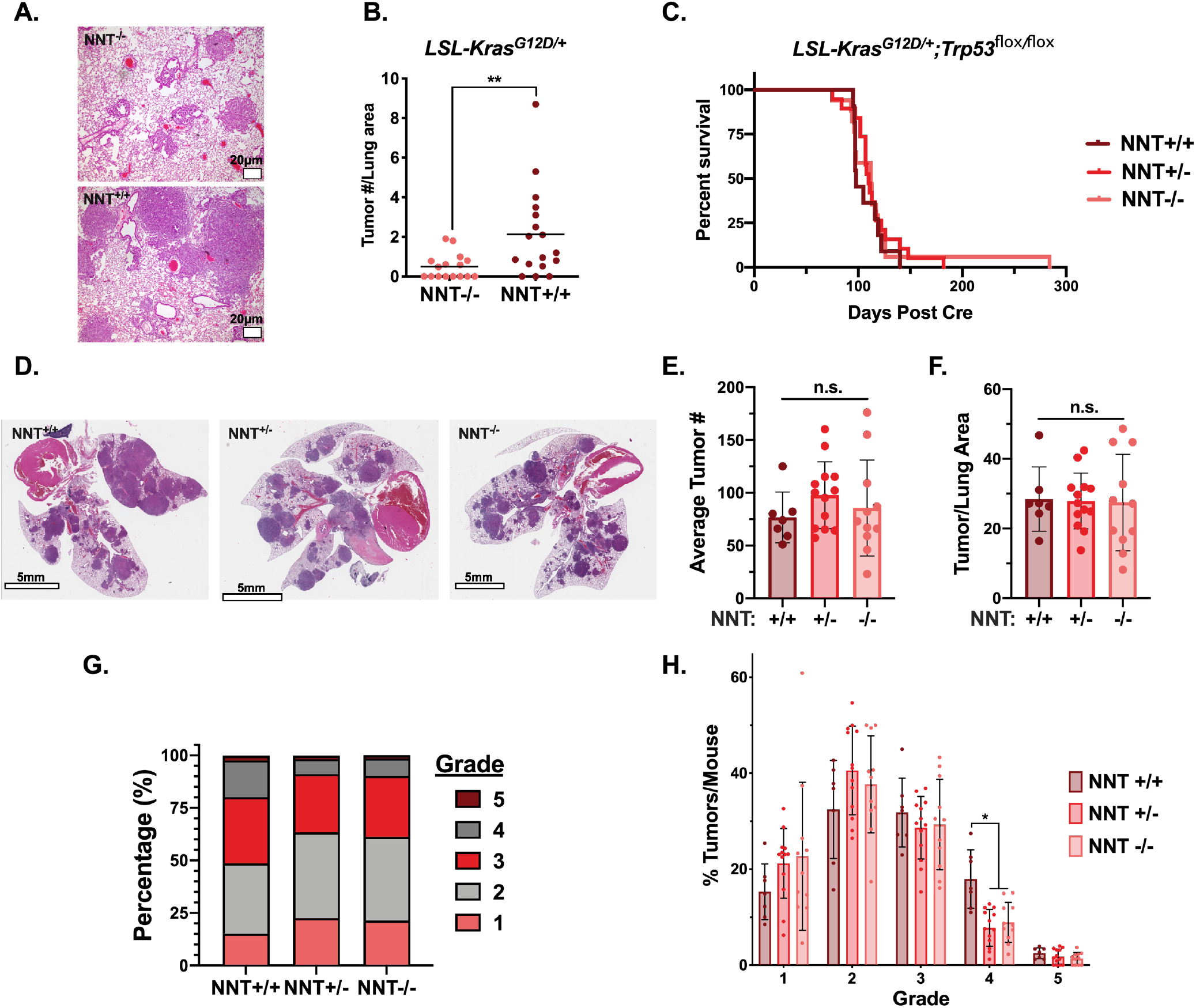
NNT supports lung tumorigenesis. **(A)** Representative hematoxylin-eosin stained lung sections of LSL-Kras^G12D/+^; NNT^−/−^ and LSL-Kras^G12D/+^; NNT^+/+^ mice 3 months after Cre induction. Bars, 20μm. **(B)** Average tumor # per lung area in LSL-Kras^G12D/+^; NNT^−/−^ (n=16) and LSL-Kras^G12D/+^; NNT^+/+^ (n=17) specimens collected at 3 months (Student’s t test). **(C)** Survival rates of LSL-Kras^G12D/+^; Trp53^flox/flox^; NNT^+/+^ (n=11), LSL-Kras^G12D/+^; Trp53^flox/flox^; NNT^+/−^ (n=19), and LSL-Kras^G12D/+^; Trp53^flox/flox^; NNT^−/−^ (n=17) following Cre induction (Log-rank test). **(D)** Representative hematoxylin-eosin stained lung sections of LSLKras^G12D/+^; Trp53^flox/flox^; NNT^+/+^ (Left), LSL-Kras^G12D/+^; Trp53^flox/flox^; NNT^+/−^ (Middle), and LSL-Kras^G12D/+^; Trp53^flox/flox^; NNT^−/−^ (Right) mice at experimental endpoint. Bars, 5mm. **(E)** Average tumor # per each lung specimen (one-way ANOVA). **(F)** Fraction of total lung that was burdened by tumor (one-way ANOVA). **(G)** Distribution of tumor grades across all tumors for Kras^G12D/+^; p53^**Δ**/**Δ**^; NNT^+/+^ (n=530 tumors), Kras^G12D/+^; p53^**Δ**/**Δ**^; NNT^+/−^ (n=1269 tumors), and Kras^G12D/+^; p53^**Δ**/**Δ**^; NNT^−/−^ (n=943 tumors). **(H)** Average frequency of each tumor grade per mouse (two-way ANOVA). For E, F, and H, LSL-Kras^G12D/+^; Trp53^flox/flox^; NNT^+/+^ (n=7), LSL-Kras^G12D/+^; Trp53^flox/flox^; NNT^+/−^ (n=13), and LSL-Kras^G12D/+^; Trp53^flox/flox^; NNT^−/−^ (n=11). For B, E, F, and H, data represented as mean ± SD. n.s., not significant; *, p<0.05; **, p<0.01.

While there were no differences in tumor burden between genotypes, we did observe differences in tumor aggressiveness as defined previously for this model (Jackson et al., 2005, DuPage et al., 2009). We found that 51.3% of tumors from NNT^+/+^ mice were of grade 3 (adenocarcinoma) or greater, whereas only 36.5% and 38.8% of tumors from NNT^+/−^ and NNT^−/−^ mice were high-grade (Fig. 1 G). This shift in tumor aggressiveness was evidenced by a significant increase in the frequency of grade 4 tumors in NNT^+/+^ mice (Fig. 1 H). Collectively, these data indicate that NNT promotes both lung tumor initiation and tumor aggressiveness.

### NNT Loss Does Not Compromise the Mitochondrial Thioredoxin Antioxidant System

To further evaluate the influence of NNT on lung tumor biology, we transitioned to human NSCLC cell lines, which exhibit varied NNT expression (Fig. S1 A). We first assessed the effect of short hairpin RNA (shRNA)-mediated knockdown of NNT on the proliferative capacity of 4 NSCLC cell lines that express NNT (A549, H1299, H2009, PC9) as well as H441 cells, which do not express NNT protein and serve as a natural negative control (Fig. 2 A). We observed that shRNA-mediated knockdown of NNT with two unique hairpins blunted the proliferative capacity of A549 and H1299 cells, while compromising the viability of H2009 and PC9 cells 5-days after lentiviral infection (Fig. 2 B). Importantly, the proliferation of H441 cells was not affected by NNT knockdown, indicating specificity of the hairpins (Fig. 2 B).

**Figure 2.**
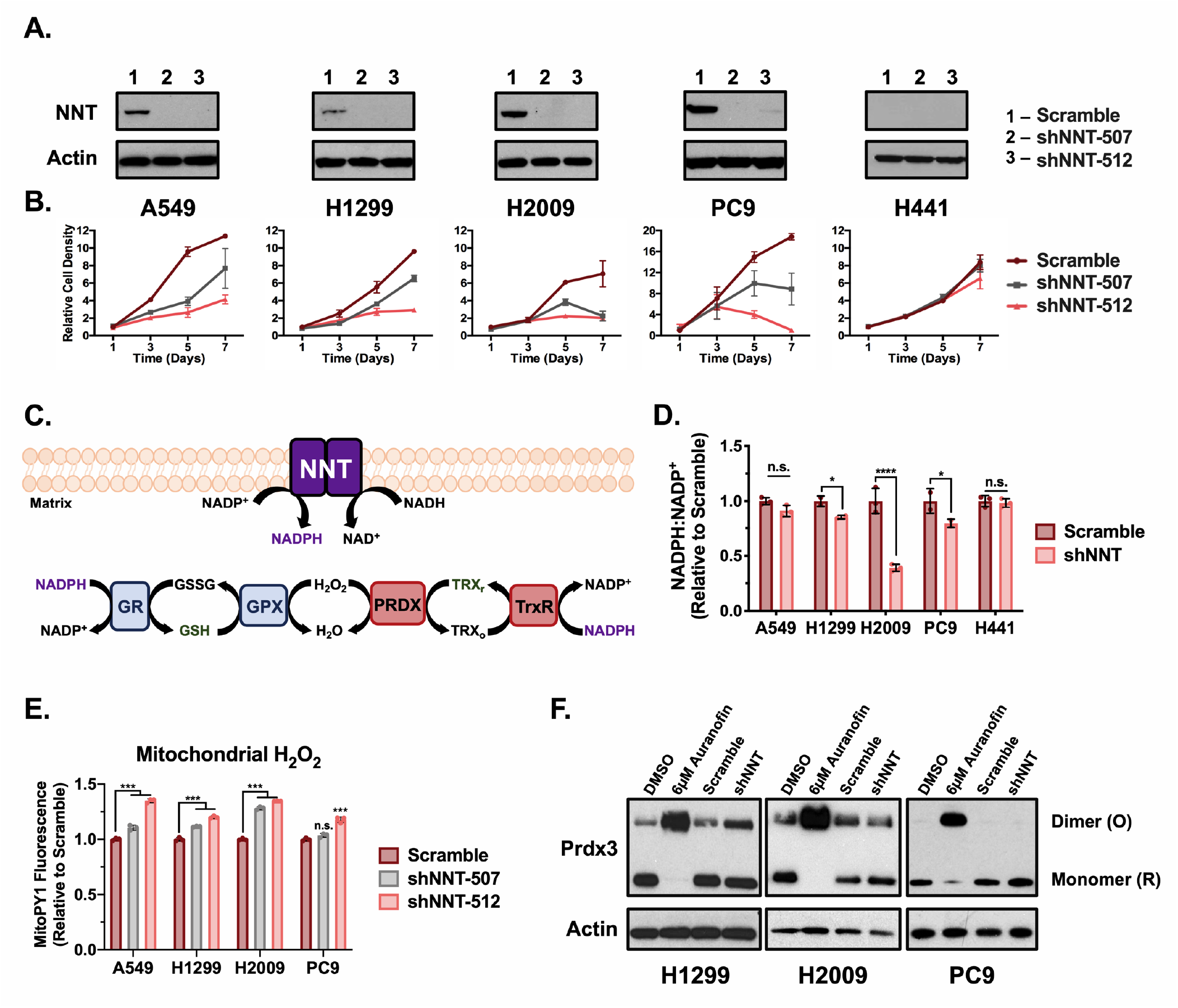
NNT loss does not compromise the mitochondrial thioredoxin antioxidant system. **(A)** Immunoblot analysis of NNT and actin (loading control) expression in NSCLC cells 3 days post-infection with scramble or shNNT lentivirus. **(B)** Plots of cell proliferation over 7 days for NSCLC cells subject to scramble or shNNT lentiviral infection. **(C)** Schematic representation of NNT’s canonical function, where NNT supplies NADPH to maintain mitochondrial antioxidant capacity. Glutathione, reduced (GSH), glutathione peroxidase (GPX), glutathione reductase (GR), glutathione, oxidized (GSSG), water (H_2_O), thioredoxin, oxidized (TRX_o_), thioredoxin, reduced (Trx_r_), thioredoxin reductase (TrxR). **(D)** NADPH:NADP+ ratio in NSCLC cells following NNT knockdown, relative to scramble infected control cells (Student’s t test). **(E)** Fluorescence of the mitochondrial H_2_O_2_ sensitive dye, MitoPY1, in NSCLC cells following NNT knockdown, relative to scramble infected control cells (one-way ANOVA). **(F)** Redox immunoblot analysis of actin (loading control) and the oxidation state of Prdx3 in parental NSCLC cells following 1-hour treatment with (1.) DMSO or (2.) 6μM auranofin or 4 days post-infection with (3.) scramble or (4.) shNNT lentivirus. Data are representative of one experiment of three experimental replicates. For B, D, and E, data are represented as mean ± SD of three technical replicates. n.s., not significant; *, p<0.05; ***, p<0.001; ****, p<0.0001.

Canonically, NNT is thought to contribute the reducing power (NADPH) required to maintain the mitochondrial glutathione and protein antioxidant systems in a reduced state (Fig. 2 C) (Rydstrom, 2006). We found that NNT knockdown reduces the cellular NADPH:NADP^+^ ratio in several cell lines (H1299, H2009, PC9) cells, while having no effect on H441 cells (Fig. 2 D).

Consistent with the notion that NNT is important for hydrogen peroxide (H_2_O_2_) detoxification, we observed modest, yet statistically significant increases in mitochondrial H_2_O_2_ levels 4-days following lentiviral infection (Fig. 2 E). We also observed modest increases in mitochondrial superoxide (•O_2_^−^) following NNT-knockdown (Fig. S1 B). However, we did not observe a discernable effect on cytosolic ROS, indicating that the impact of NNT on ROS accumulation is largely confined to the mitochondria (Fig. S1 C).

To determine if these increases in mitochondrial ROS species related to the mitochondrial antioxidant systems, we assessed the oxidation state of the mitochondrial peroxiredoxin (Prdx3) through redox western blotting. H_2_O_2_ detoxification by Prdx3 induces dimerization of the protein through a pair of cysteine disulfide bonds that must be reduced by thioredoxin in order to restore Prdx3 antioxidant function. Thus, accumulation of dimerized Prdx3 is an indication of Prdx3 oxidation and a surrogate marker for mitochondrial oxidative stress (Cox et al., 2009). While we found that treatment with auranofin, an inhibitor of thioredoxin reductase resulted in substantial oxidation of Prdx3, the loss of NNT did not increase Prdx3 oxidation relative to scramble-infected control cells (Fig. 2 F). Furthermore, NNT knockdown did not sensitize NSCLC cells to treatment with tert-butyl hydroperoxide (tbHP), cumene hydroperoxide (CHP), or the mitochondrial-targeted menadione (Figs. S1 D, E, and F). Collectively, these results indicate that NNT is important for the proliferative capacity of NSCLC cells but that loss of NNT activity does not affect the thioredoxin antioxidant system or elicit significant mitochondrial oxidative stress.

### NNT Loss Compromises Mitochondrial Oxidative Capacity

Given the localization of NNT within the inner mitochondrial membrane and its ability to influence proton flux across the membrane and reducing power, we sought to interrogate whether NNT was important for mitochondrial oxidative metabolism. First, we employed the Seahorse extracellular flux analyzer to perform a MitoStress test as an assessment of the impact of NNT loss on general mitochondrial oxidative function. We observed that the oxygen consumption rate (OCR) of NNT-deficient cells was reduced relative to scramble control cells in response to the sequential delivery of mitochondrial inhibitors (Fig. 3 A). Notably, the maximal respiratory capacity of NNT-deficient cells was significantly lower independent of an effect on uncoupled respiration (Figs. 3 B and S2 A). This is indicative of a mitochondrial oxidative defect that is unrelated to NNT’s influence on the proton gradient.

**Figure 3.**
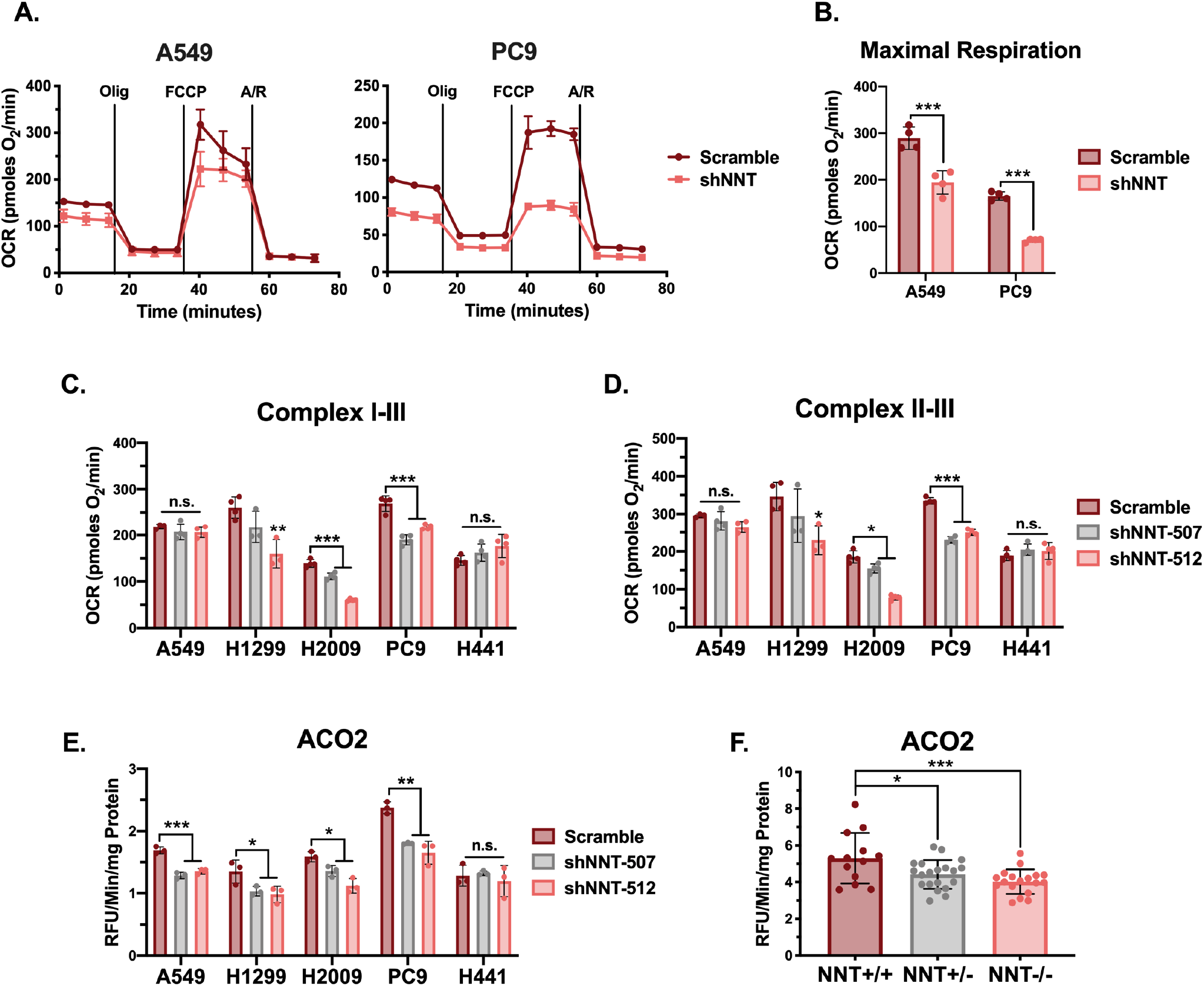
NNT loss compromises mitochondrial oxidative capacity. **(A)** Plots of OCR in A549 and PC9 cells subject to NNT knockdown. Cells were supplemented with 10mM glucose and 1mM glutamine and then sequentially challenged with 1μM oligomycin (Olig), 1μM (A549) or 0.5μM (PC9) of carbonyl cyanide 4-(trifluoromethoxy)phenylhydrazone (FCCP), and 1μM each of antimycin A (A) and rotenone (R). **(B)** Average maximal respiratory capacity of A549 and PC9 cells following infection with either scramble or shNNT lentivirus (Student’s t test). **(C)** Average complex I-III activity following stimulation with 10mM pyruvate and 1mM malate in NSCLC cells subject to NNT knockdown (one-way ANOVA). **(D)** Average complex II-III activity following stimulation with 10mM succinate in NSCLC cells subject to NNT knockdown (one-way ANOVA). **(E)** Average ACO2 activity in mitochondrial lysates of NSCLC cells following NNT knockdown (one-way ANOVA). **(F)** Average ACO2 activity in mitochondrial lysates of lung tumors collected from LSL-Kras^G12D/+^; Trp53^flox/flox^; NNT^+/+^ (n=13 tumors from 5 mice), LSL-Kras^G12D/+^; Trp53^flox/flox^; NNT^+/−^ (n=20 tumors from 5 mice), and LSL-Kras^G12D/+^; Trp53^flox/flox^; NNT^−/−^ (n=18 tumors from 4 mice) mice (one-way ANOVA). For A-E, data are representative of one experiment of three experimental replicates. For A-E, data are represented as mean ± SD of at least three technical replicates. n.s., not significant; *, p<0.05; **, p<0.01; ***, p<0.001.

Mitochondrial oxidative metabolism is dependent on a functional ETC, which consists of protein complexes with resident Fe-S clusters that mediate electron transport. Considering that NADPH is required for Fe-S cluster biosynthesis, we endeavored to examine if the decrease in mitochondrial respiratory function following NNT knockdown was linked to the Fe-S proteins within the ETC (Webert et al., 2014). While the MitoStress test allows for a general analysis of respiratory function, it does not allow for the evaluation of individual ETC complexes. Therefore, we performed a more specialized Seahorse-based protocol that permits the sequential analysis of the Fe-S cluster-dependent respiratory complexes (I, II, III) (Salabei et al., 2014). We found that in response to feeding with the complex I substrates, pyruvate and malate, the activity of complex I-III was significantly reduced following NNT knockdown (Fig. 3 C). Furthermore, NNT-deficient cells also exhibited significantly reduced OCR in response to succinate, indicative of reduced complex II-III activity (Fig. 3 D). As expected, NNT knockdown did not alter the Fe-S cluster-dependent respiratory chain activities of H441 cells (Figs. 3 C and D).

In addition to sustaining electron flux through the ETC, Fe-S clusters also suport the enzymatic function of other proteins critical to oxidative metabolism. To determine if NNT contributes to the function of other Fe-S proteins, we assessed the activity of aconitase (ACO2), a Fe-S protein of the TCA cycle. We found that NNT knockdown significantly reduced ACO2 activity in those NSCLC lines with NNT expression (Fig. 3 E). This reduction in ACO2 activity occurred independently from a decrease in ACO2 expression, suggesting that the change in activity was due to a functional deficit (Fig S2 B).

Diminished ACO2 activity is likely to disrupt TCA cycling, leading to a reduced capacity to generate the reducing equivalents needed to drive ETC flux. To ensure that the decreased respiratory chain phenotypes we observed were not simply consequences of this ACO2 defect, we assessed respiratory chain function in response to glutamate and malate stimulation. Glutamate carbon can enter the TCA cycle as α-ketoglutarate, permitting us to circumvent the need for ACO2, which is required for the initial turn of pyruvate carbon through the cycle. Regardless, NNT-deficient cells exhibited equally disrupted respiratory chain function in response to glutamate as they did to pyruvate (Fig S2 C and D). To supplement the analyses of ACO2 function in out NSCLC cell lines, we also evaluated the influence of NNT expression on ACO2 activity in KP lung tumors. We found that ACO2 activity was significantly higher in tumors from NNT+/+ mice than those of NNT+/− or NNT−/−, with tumors lacking NNT exhibiting the lowest activity (Fig. 3 F). Like in the human cell lines, this difference in activity was not the result of differential ACO2 expression (Fig. S2 E).

### An Exogenous Source of NADPH Sustains Fe-S Protein Function Following NNT Loss

To determine if the decreases in mitochondrial Fe-S protein function associated with NNT knockdown were related to the accompanying decrease in NADPH:NADP^+^, we sought to provide an exogenous source of mitochondrial NADPH. To achieve this, we chose the yeast mitochondrial NADH kinase, pos5p, which phosphorylates NADH to yield NADPH (Pain et al., 2010). Though pos5p has been exogenously expressed in bacteria previously (Lee et al., 2013), it has never been introduced into a mammalian system. To monitor if we could efficiently express pos5p protein in the mitochondria of our human NSCLC cells, we modified the yeast protein to include an HA-tag. Western blot analysis of fractionated lysates revealed successful expression of HA-tagged pos5p in the mitochondria of H1299, H2009, and PC9 cells (Fig. 4 A). These lines were chosen to evaluate the ability of pos5p to rescue the Fe-S protein defects associated with NNT loss as they exhibited the most severe response to NNT knockdown. Importantly, we did not observe any adverse effects of pos5p expression on mitochondrial function in our NSCLC cells (Fig. S3).

**Figure 4.**
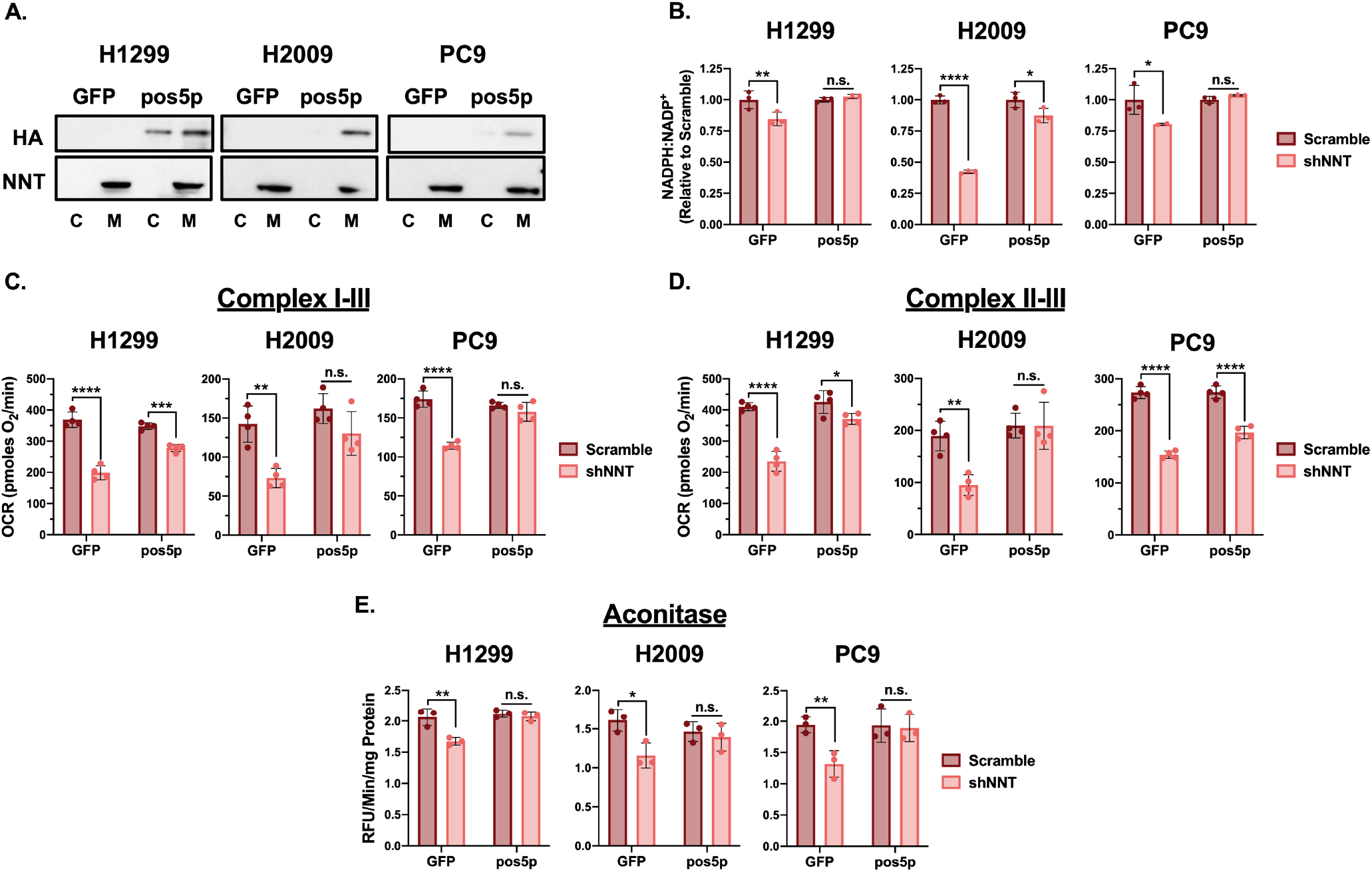
An exogenous source of NADPH sustains Fe-S protein function following NNT loss. **(A)** Immunoblot analysis of HA-tag and NNT (mitochondrial control) expression in cytosolic and mitochondrial lysates taken from GFP or pos5p expressing NSCLC cells. **(B)** NADPH:NADP+ ratio in GFP or pos5p expressing NSCLC cells following NNT knockdown, relative to scramble infected control cells (two-way ANOVA). **(C)** Average complex I-III activity following stimulation with 10mM pyruvate and 1mM malate in GFP or pos5p expressing NSCLC cells subject to NNT knockdown (two-way ANOVA). **(D)** Average complex II-III activity following stimulation with 10mM succinate in GFP or pos5p expressing NSCLC cells subject to NNT knockdown (two-way ANOVA). **(E)** Average ACO2 activity in mitochondrial lysates of GFP or pos5p expressing NSCLC cells following NNT knockdown (two-way ANOVA). Data are representative of one experiment of three experimental replicates. For B-E, data are represented as mean ± SD of at least three technical replicates. n.s., not significant; *, p<0.05; **, p<0.01; ***, p<0.001; ****, p<0.0001.

Expression of pos5p rescued the decrease in the cellular NADPH:NADP^+^ ratio elicited by NNT knockdown (Fig. 4 B). This corresponded with an attenuation of the decrease in respiratory chain complex activities following knockdown of those cells expressing pos5p (Figs. 4 C and D). Moreover, pos5p expression fully rescued the decrease in ACO2 activity associated with NNT knockdown (Fig. 4 E). Together, these data indicate that maintaining NADPH levels upon loss of NNT expression protects Fe-S protein function in NSCLC cells.

### NNT Loss Does Not Disrupt Fe-S Cluster Biosynthesis

Given that an exogenous source of mitochondrial NADPH attenuated the Fe-S protein defects associated with NNT knockdown and that NADPH is required for efficient and sustained Fe-S cluster biosynthesis, we next sought to determine if NNT activity sustained this process. Fe-S cluster biosynthesis occurs at a multiprotein complex consisting in part of the cysteine desulfurase (NFS1) and iron-sulfur scaffold protein (ISCU) (Johnson et al., 2005). Loss of either compromises cluster biosynthesis and is associated with mitochondrial defects (Fosset et al., 2006, Crooks et al., 2018). Therefore, we introduced shRNAs targeting either NFS1 or ISCU (Alvarez et al., 2017) to establish the effects of disrupting Fe-S cluster biosynthesis on the respiratory chain and ACO2.

NFS1 knockdown significantly diminished the activities of the respiratory chain complexes in response to both pyruvate/malate as well as succinate in H2009 cells, whereas only complex II-III activity was significantly reduced in PC9 cells (Fig. 5 A and B). Alternatively, loss of ISCU expression significantly blunted OCR in response to pyruvate/malate and succinate in both cell lines (Fig. 5 A and B). Furthermore, knockdown of either NFS1 or ISCU significantly reduced ACO2 activity across cell lines (Fig. 5 C). Intriguingly, the deficits elicited by NNT knockdown were of equal magnitude to those resultant from targeting these bona fide components of the Fe-S cluster biosynthetic machinery (Figs. 5 A, B, and C).

**Figure 5.**
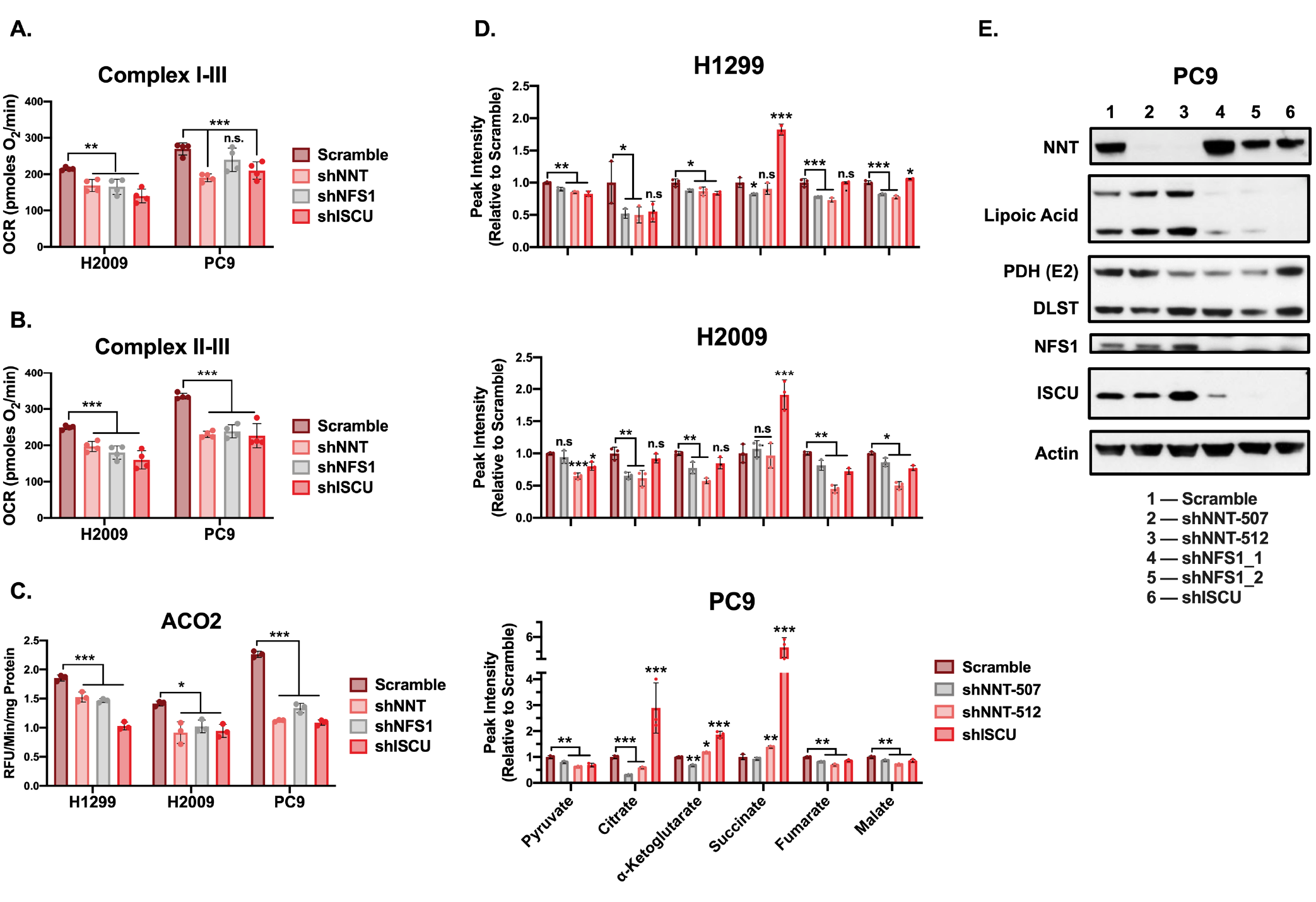
NNT loss does not disrupt Fe-S cluster biosynthesis. **(A)** Average complex I-III activity following stimulation with 10mM pyruvate and 1mM malate in H2009 and PC9 cells subject to NNT, NFS1, or ISCU knockdown (one-way ANOVA). **(B)** Average complex II-III activity following stimulation with 10mM pyruvate and 1mM malate in H2009 and PC9 cells subject to NNT, NFS1, or ISCU knockdown (one-way ANOVA). **(C)** Average ACO2 activity in mitochondrial lysates of NSCLC cells following NNT, NFS1, or ISCU knockdown (one-way ANOVA). **(D)** Relative abundance of TCA cycle intermediates in extracts of NSCLC cells subject to NNT or ISCU knockdown (one-way ANOVA). **(E)** Immunoblot analysis of NNT, lipoic acid, PDH-E2, DLST, NFS1, ISCU, and actin (loading control) expression in PC9 cells following infection with (1) scramble, (2) shNNT-507, (3) shNNT-512, (4) shNFS1_1, (5) shNFS1_2, or (6) ISCU lentivirus. For A-C, data are representative of one experiment of three experimental replicates. For A-C, data are represented as mean ± SD of at least three technical replicates. For D, data are represented as mean ± SD of three biological replicates. n.s., not significant; *, p<0.05; **, p<0.01; ***, p<0.001.

To demonstrate that the Fe-S protein defects we observed have a functional impact on the mitochondrial metabolism of these NSCLC cells, we performed liquid chromatography-high resolution mass spectrometry (LC-HRMS)-based metabolomics on cells subjected to NNT or ISCU knockdown. Analysis of TCA cycle metabolites from these cells revealed significant alterations in the abundance of most intermediates across cell lines (Fig. 5 D). These are indicative of a severe disruption of oxidative metabolism and consistent with the described defects in Fe-S protein function. Specifically, we observed a depletion of pyruvate, malate and fumarate following disruption of both NNT and ISCU expression. While citrate levels were depleted in NNT-deficient cells, there was no consistent effect of ISCU knockdown. Alternatively, ISCU-deficient cells exhibited a striking accumulation of succinate that was absent in NNT knockdown cells (Fig. 5 D).

Though ACO2 is the only Fe-S protein in the TCA cycle, several other TCA cycle proteins depend on the function of an additional mitochondrial Fe-S protein, lipoic acid synthetase (LIAS). LIAS is required for lipoic acid synthesis and the eventual conjugation of crucial lipoate moieties to components of PDH (E2) and α-ketoglutarate dehydrogenase (dihydrolipoamide S-succinyltransferase, DLST), among others. LIAS is critically sensitive to disruptions in Fe-S cluster biosynthesis, as its resident Fe-S cluster is consumed during catalysis, imposing a requirement for continual cluster turnover (Crooks et al., 2018). Indeed, disruption of NFS1 and ISCU expression resulted in a substantial reduction in PDH-E2 and DLST lipoylation in PC9 cells (Fig. 5 E). However, NNT knockdown had no effect on protein lipoylation (Fig. 5 E). Collectively, these data suggest that while NNT elicits enzymatic and metabolic defects reminiscent of those associated with the disruption of Fe-S cluster biosynthesis, it is unlikely that NNT directly influences this process.

### NNT Loss Disrupts Fatty Acid Metabolism

In addition to the depletion of TCA cycle intermediates, LC-HRMS analysis of NNT-deficient cells revealed a metabolic signature indicative of dysregulated fatty acid metabolism. NNT knockdown promoted the accumulation of both saturated and unsaturated fatty acids relative to scramble-infected controls (Fig 6 A). This was accompanied by a significant accumulation of the β-oxidation substrate, palmitoylcarnitine, in NNT-deficient cells (Fig. 6 B). Given the established respiratory defects seen in NNT-deficient cells, we anticipated that the increase in palmitoylcarnitine was a result of decreased fatty acid oxidation. Consistent with this, we found that OCR linked to palmitate oxidation was reduced in H1299 and PC9 cells following NNT knockdown (Fig 6 C). To evaluate if the accumulation of fatty acids following NNT knockdown is a potential liability, we challenged NNT-deficient cells with the saturated fatty acid palmitate for 24-hours. We found that NNT knockdown sensitized H1299 and H2009 cells to palmitate treatment (Fig. 6 D). Furthermore, NNT knockdown significantly sensitized H1299 and PC9 cells to treatment with the monounsaturated fatty acid, oleate (Fig. 6 E). These data suggest that NNT loss perturbs fatty acid metabolism in NSCLC cells that may serve as an exploitable vulnerability.

**Figure 6.**
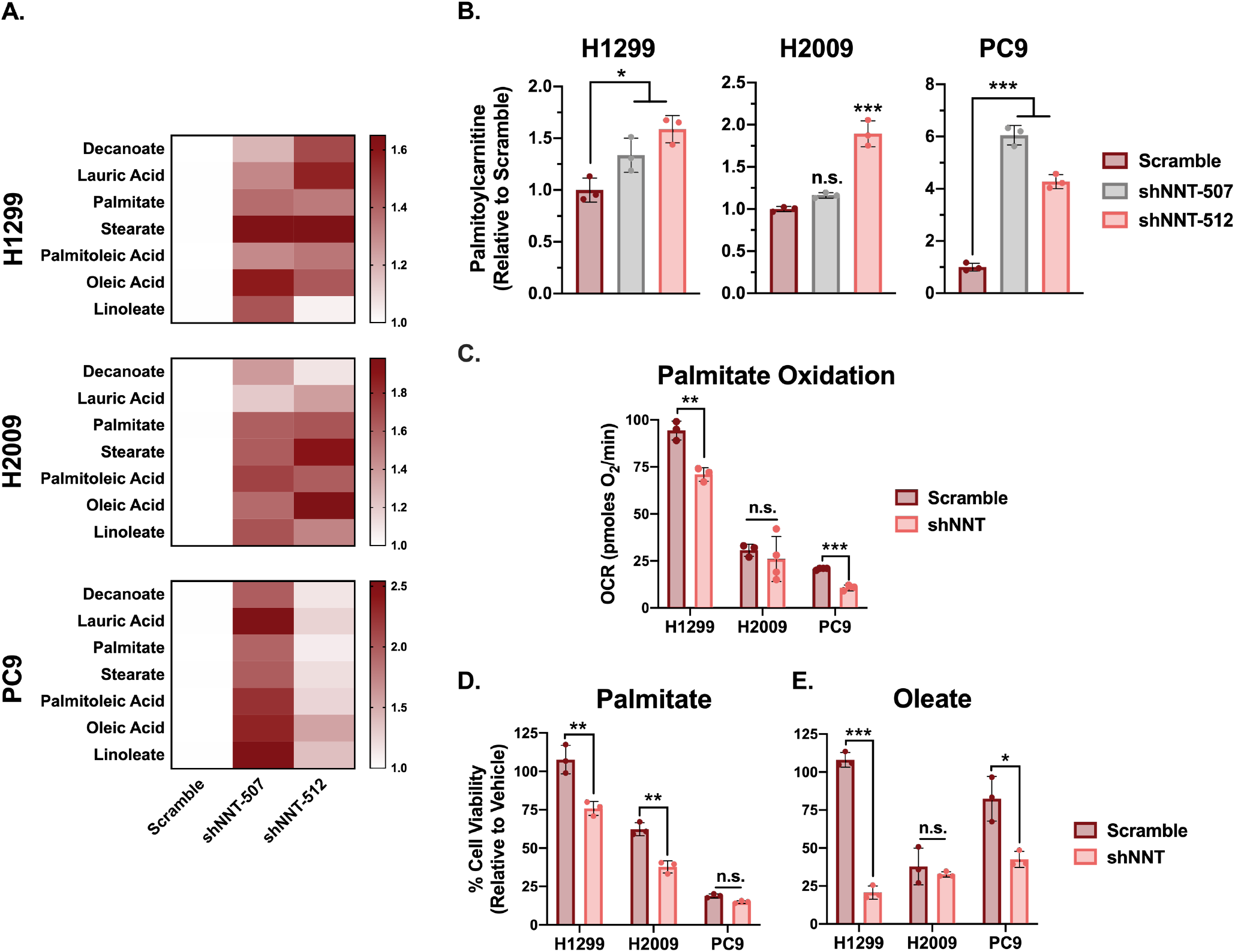
NNT loss disrupts fatty acid metabolism. **(A)** Heat map representation of the relative abundances of saturated (decanoate, lauric acid, palmitate, stearate) and unsaturated (palmitoleic acid, elaidic acid, linoleate) fatty acids in extracts of NSCLC cells following NNT knockdown. **(B)** Relative abundance of palmitoylcarnitine in extracts of NSCLC cells subject to NNT knockdown (one-way ANOVA). **(C)** Measures of OCR coupled to the oxidation of exogenous palmitate-BSA in NSCLC cells subject to NNT knockdown (Student’s t test). **(D-E)** Viability of NSCLC cells subject to scramble or shNNT lentiviral infection following 24-hour treatment with (D) 200μM of palmitate or (E) 400μM of oleate (Student’s t test). Cell viability was determined relative to BSA treated controls. For C-E, data are representative of one experiment of three experimental replicates. For A, data are represented as the mean fold increase relative to scramble of three biological replicates. For B, data are represented as mean ± SD of three biological replicates. For C-E, data are represented as mean ± SD of at least three technical replicates. n.s., not significant; *, p<0.05; **, p<0.01; ***, p<0.001.

### Mitochondrially-Targeted Catalase Rescues Fe-S Protein Function Following NNT Loss

Fe-S clusters are exquisitely sensitive to oxidation by molecular oxygen and more deleterious species (Flint et al., 1993, Djaman et al., 2004, Alvarez et al., 2017). Though we did not observe changes in the oxidation state of the mitochondrial protein antioxidant system, that does not preclude that the modest induction of mitochondrial ROS following NNT knockdown is sufficient to oxidize these sensitive cofactors. To interrogate this possibility, we employed a mitochondrially-targeted catalase (MitoCatalase) to enhance mitochondrial antioxidant capacity. We successfully overexpressed MitoCatalase in H1299, H2009, and PC9 cells (Fig. 7 A). These cells exhibited an enhanced capacity to clear mitochondrial H_2_O_2_ upon challenge with 100μM of exogenous H_2_O_2_, (Fig. S4 A). Furthermore, these MitoCatalase expressing cells were more resistant to menadione treatment, indicating functionality within the mitochondria (Fig. S4 B).

**Figure 7.**
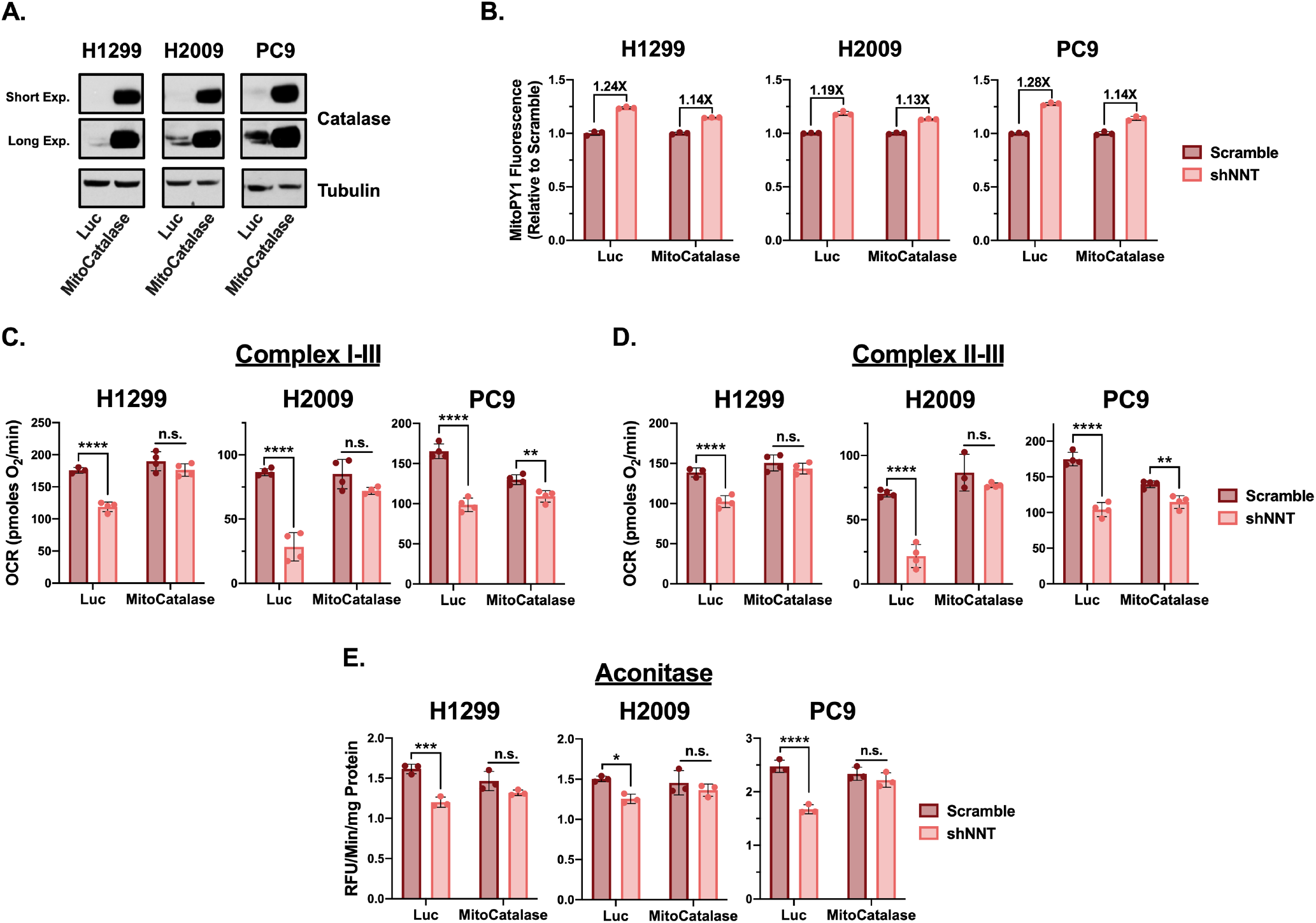
Mitochondrially targeted catalase rescues Fe-S protein function following NNT loss. **(A)** Immunoblot analysis of catalase and tubulin (loading control) expression in (1) Luciferase (Luc) or (2) MitoCatalase expressing NSCLC cells. **(B)** Fold inductions of MitoPY1 fluorescence in shNNT-infected Luc or MitoCatalase expressing NSCLC cells. **(C)** Average complex I-III activity following stimulation with 10mM pyruvate and 1mM malate in Luc or MitoCatalase expressing NSCLC cells subject to NNT knockdown (two-way ANOVA). **(D)** Average complex II-III activity following stimulation with 10mM succinate in Luc or MitoCatalase expressing NSCLC cells subject to NNT knockdown (two-way ANOVA). **(E)** Average ACO2 activity in mitochondrial lysates of Luc or MitoCatalase expressing NSCLC cells following NNT knockdown (two-way ANOVA). Data are representative of one experiment of three experimental replicates. For B-E, data are represented as mean ± SD of at least three technical replicates. n.s., not significant; *, p<0.05; **, p<0.01; ***, p<0.001; ****, p<0.0001.

The expression of MitoCatalase also partially attenuated the modest induction of mitochondrial H_2_O_2_ associated with NNT knockdown. This corresponded with an attenuation of the reduction in respiratory chain complex activities following NNT knockdown (Fig. 7 C and D). Moreover, MitoCatalase expression rescued ACO2 activity in response to NNT knockdown (Fig. 7 E). This rescue of ACO2 activity was specific to NNT, as NFS1 deficient cells exhibited reduced ACO2 activity even in the presence of MitoCatalase (Fig. S4 C). Collectively, these data indicate that enhancing the mitochondrial capacity to detoxify H_2_O_2_ protects against the Fe-S protein deficits associated with NNT knockdown, further arguing against a connection between NNT and Fe-S cluster biosynthesis.

## Discussion

We demonstrate here for the first time that NNT expression supports lung tumorigenesis in two genetically engineered mouse models (GEMM) of NSCLC. These GEMMs were crossed with the natural deletion variant of NNT from a C57BL/6J background (Toye et al., 2005). Intriguingly, C57BL/6J mice are largely resistant to tumor formation (Law et al., 1967), which in light of our findings, suggests a potential role for NNT-deficiency in this phenotype. Our observation that NNT expression significantly enhances Kras^G12D^-driven lung tumor formation (Figs. 1A and B) is consistent with previous work with this GEMM demonstrating a need for proteins involved in mitochondrial redox metabolism (Weinberg et al., 2010, DeNicola et al., 2011, Davidson et al., 2016, Mayers et al., 2016, Padanad et al., 2016, Rao et al., 2019). A requirement for mitochondrial function in this model aligns with the robust mitochondrial metabolism exhibited by human lung tumors (Hensley et al., 2016).

The oxidation of carbon within the mitochondria stimulates ETC flux, which precipitates the formation of superoxide and its subsequent conversion to other ROS species. This necessitates a functional antioxidant system to prevent an unsustainable accumulation of macromolecular oxidation that can compromise ETC function and mitochondrial integrity. Indeed, activation of nuclear factor erythroid 2-related factor 2 (Nrf2), the master regulator of cellular antioxidant capacity, is a common feature of NSCLC (Singh et al., 2006) and enhances lung tumorigenesis (DeNicola et al., 2011, Romero et al., 2017). Yet, in order to sustain this increased antioxidant capacity, cells must maintain a reduced NADPH:NADP^+^ ratio, especially in the mitochondria (Klingenberg and Slenczka, 1959). Given the established function of NNT in sustaining mitochondrial antioxidant capacity through regulation of the NADPH:NADP^+^ ratio (Lopert and Patel, 2014, Ronchi et al., 2016), we anticipated that NNT would serve a similar role in contributing to mitochondrial function in NSCLC. Surprisingly, while we did determine that NNT contributes to the NADPH:NADP+ ratio in NSCLC cells, the accompanying increase in mitochondrial ROS was not sufficient to compromise the mitochondrial protein antioxidant system, nor sensitize NNT-deficient cells to treatment with oxidants (Figs. 2 and S1). This is in direct contrast to what was observed in a model of adrenal adenocarcinoma (Chortis et al., 2018). Though, it was previously established that NNT regulation of global oxidation is critical to adrenal physiology and NNT deficiency manifests with adrenal insufficiency in patients (Roucher-Boulez et al., 2016, Meimaridou et al., 2018). This indicates that while NNT serves to supplement the NADPH pool across tissue types (Lopert and Patel, 2014, Fisher-Wellman et al., 2015, Ronchi et al., 2016, Meimaridou et al., 2018), its functional contribution to redox homoeostasis may vary, especially with regards to malignancy (Gameiro et al., 2013, Ho et al., 2017, Chortis et al., 2018, Li et al., 2018).

It has been shown in disparate cancer cell lines that disruption of redox homeostasis following the loss of NNT expression induces metabolic rewiring, marked by changes in fuel utility (Gameiro et al., 2013, Ho et al., 2017). In an endothelial cell line derived from the ascitic fluid of a patient with liver adenocarcinoma, NNT knockdown was associated with reduced flux of both glucose and glutamine carbon through the TCA cycle coupled with a shift towards reductive glutamine metabolism (Ho et al., 2017). This is in contrast to melanoma cells, which exhibit increased glucose flux through the TCA cycle at the expense of reductive glutamine metabolism (Gameiro et al., 2013). This finding aligns with a study by Mullen et al., which demonstrated a connection between NNT-derived NADPH and reductive glutamine metabolism (Mullen et al., 2014). Though we did not assess carbon flux through the TCA cycle, our data indicate a reduced capacity to oxidize either glucose or glutamine carbon in NSCLC cells following NNT knockdown (Figs. 3 and S2). The divergence in metabolic consequences following disruption of NNT expression in various cancers further suggests a context dependency to its function.

Through our LC/MS-based assessment of the impact of NNT loss on NSCLC metabolism, we revealed a dysregulation of mitochondrial metabolism marked by a severe depletion of most TCA cycle intermediates (Fig. 5 D). This corroborated our earlier findings that NNT knockdown diminished the function of Fe-S proteins critical to mitochondrial oxidative metabolism. While the impact of NNT loss on Fe-S protein function was strikingly similar to that of disrupting the Fe-S cluster biosynthetic machinery, there were distinct effects on the TCA cycle that distinguished NNT loss from ISCU knockdown (Fig. 5). This deviation is potentially attributable to the influence of ISCU but not NNT on protein lipoylation. Unlike the clusters associated with ACO2 or the respiratory chain, the 4Fe-4S cluster that mediates LIAS catalysis is consumed in the process to contribute sulfur for synthesis of lipoic acid (Parry and Trainor, 1978, Miller et al., 2000, Crooks et al., 2018). This activity permits lipoylation of components of PDH and α-ketoglutarate dehydrogenase, which are required for TCA cycling. The lack of a discernable effect on the lipoylation status of PDH and DLST following NNT knockdown indicates that NNT does not influence these proteins in the same manner as ISCU (Fig. 5 E). Still, the metabolic effects of NNT loss were very reminiscent of those elicited by the acute loss of Fe-S clusters, which included an accumulation of fatty acids (Crooks et al., 2018). In that context, an ISCU deficiency promoted de novo fatty acid synthesis from citrate carbon that could not be efficiently oxidized in the TCA cycle due to the corresponding loss of Fe-S protein function. It is unlikely that NNT loss promotes fatty acid synthesis in NSCLC cells due to the significant demand for NADPH in the generation of fatty acids and the observed decrease in NADPH availability associated with NNT knockdown (Fig 2 D). Rather, this is likely a result of increased uptake, which occurs in NSCLC cells incapable of de novo synthesis (Migita et al., 2008). This blockade of fatty acid synthesis elicited an enhanced sensitivity to exogenous palmitate treatment, which we also observed in our NNT-deficient cells (Fig. 6 D). Regardless, fatty acid accumulation is a shared response to the disruption of either NNT or ISCU expression.

The differential effects of NNT loss on mitochondrial Fe-S proteins with respect to the catalytic function of the resident cluster(s) suggested that NNT may influence cluster integrity rather than availability. Fe-S clusters are exquisitely sensitive to oxidation, including by molecular oxygen (Crack et al., 2014, Alvarez et al., 2017). Cluster oxidation is associated with dislocation of an iron atom that renders the cluster inactive (Djaman et al., 2004). The highly oxygenated pulmonary environment likely dictates the substantial mitochondrial oxidation exhibited by human lung tumors (Hensley et al., 2016), as oxidative phosphorylation exploits the detoxification of molecular oxygen for an energetic benefit. Interestingly, NFS1 is positively selected for in NSCLC and is shown to support sustained Fe-S cluster biosynthesis to protect against the excessive oxygen challenge associated with residency in the lung (Alvarez et al., 2017). These clusters are also subject to oxidation by the reactive products of the same metabolism that protects against the accumulation of molecular oxygen (Flint et al., 1993, Djaman et al., 2004). Though we do not observe a substantial induction of ROS associated with NNT knockdown (Figs. 2 E and S1 B), we are able to rescue the effects of NNT loss on Fe-S protein function with MitoCatalase (Fig. 7). Moreover, the differential effect of NNT and ISCU loss on LIAS function may further reflect the importance of NNT to cluster integrity rather than synthesis. Given that the cluster associated with LIAS must be consistently turned over due to its consumption during catalysis, it is likely less prone to oxidation.

Considering our similar ability to rescue Fe-S protein function with pos5p (Fig. 5), our collective data indicate a potential role for NNT-derived NADPH in the mitigation of Fe-S cluster oxidation. This is intuitive considering that NNT is localized to the same membrane as the respiratory chain, which is a significant source of mitochondrial ROS. Thus, NNT may provide a regionalized source of reducing power to protect the integrity of the ETC. With regards to the potential impact of NNT activity on the soluble ACO2, there is substantial evidence that ACO2 physically associates with TCA cycle enzymes to form dynamic assemblies to enhance substrate channeling (Porpaczy et al., 1987, Morgunov and Srere, 1998, Fernie et al., 2018). Moreover, several of the NADH yielding dehydrogenases associate with the IMM, likely enhancing oxidation by complex (D’Souza and Srere, 1983, Sumegi and Srere, 1984a, Sumegi and Srere, 1984b). Together, this suggests a likely spatial association between the TCA cycle machinery and the IMM permitting regulation by NNT. Indeed, there is existing evidence that NNT contributes to a redox cycle with PDH (Fisher-Wellman et al., 2015). As described, NNT consumes the NADH yielded by PDH to generate the NADPH required to support the detoxification of H_2_O_2_ that is also produced as a byproduct of PDH catalysis (Fisher-Wellman et al., 2013). This NNT-dependent redox circuit was linked to respiratory capacity and energy expenditure in mice (Fisher-Wellman et al., 2015).

Altogether, our study demonstrates that NNT is of significance to lung tumor biology, in part through the regulation of Fe-S proteins that facilitate mitochondrial metabolism. In contrast to previous studies evaluating NNT function, we describe a nuanced influence on mitochondrial redox homeostasis in NSCLC, where NNT activity likely mitigates regionalized oxidative stress stemming from the substantial oxidative metabolism exhibited by lung tumors. Our findings further indicate a necessity for mitochondrial metabolism in lung tumorigenesis, underscoring the therapeutic potential of augmenting mitochondrial function (Weinberg and Chandel, 2015).

## Materials and Methods

### Mice

*LSL-Kras*^*G12D*^ mice (Jackson et al., 2001) were crossed with NNT-deficient C57BL/6J mice and interbred to generate *LSL-Kras*^*G12D*^; *NNT*^−/−^ and *Kras^G12D^*; *NNT*^+/+^ mice for lung tumor studies. *LSL-Kras*^*G12D*^ and *Trp53*^*flox*^ mice (Jackson et al., 2005) on a C57BL/6J and 129 mixed background were interbred to generate *LSL-Kras*^*G12D*^; *Trp53*^*flox/flox*^; *NNT*^−/−^, *LSL-Kras*^*G12D*^; *Trp53*^*flox/flox*^; *NNT*^+/−^, and *LSL-Kras*^*G12D*^; *Trp53*^*flox/flox*^; *NNT*^+/+^ mice for lung tumor studies. To induce lung tumor formation, mice under isofluorane anesthesia were infected intranasally with 10^7^ PFU of adenoviral-Cre (University of Iowa) as previously described (Jackson et al., 2001). All mice studies were approved by and conducted in accordance to the ethical standards established by the University of South Florida IACUC (protocol # R IS00003893).

### Tumor Analysis

For analyses of Kras^G12D^; NNT^−/−^ and Kras^G12D^; NNT^+/+^ tumors, mice were euthanized 3 months following tumor induction with adenoviral-Cre. Lungs were collected and fixed in 10% formalin overnight and then embedded in paraffin for subsequent sectioning. Sections were deparafinized in xylene and then rehydrated in a graded series of ethanol solutions. Slides were then sequentially stained with hematoxylin and eosin, dehydrated in ethanol and xylene, coverslipped and then dried overnight. Slides were scanned with the Aperio imager (Leica Biosystems) and each lung specimen was analyzed with ImageScope (Aperio). For analyses of Kras^G12D^; p53^**Δ**/**Δ**^; NNT^−/−^, Kras^G12D^; p53^**Δ**/**Δ**^; NNT^+/−^, and Kras^G12D^; p53^**Δ**/**Δ**^; NNT^+/+^ tumors, mice were enrolled in a survival study and allowed to reach an experimental endpoint in agreement with ethical standards. Lung specimens were collected, H&E slides were generated, and histopathology analysis was performed to grade lesions on a (1-5) scale based on criteria previously established for this model (Jackson et al., 2005, DuPage et al., 2009).

### Lentiviral Generation and Infection

Lenti-X 293 T cells (Clontech) at 90% confluency were co-transfected with the plasmid of interest and the packaging plasmids pCMV-dR8.2 dvpr (addgene #8455) and pCMV-VSV-G (addgene #8454) in the presence of JetPRIME (Polyplus). Cells were infected with lentivirus and 8μg/mL polybrene for 6-hours at an optimized dilution established with the Lenti-X™ Gostix™ Plus system (Takara).

### Cell Lines and Culture

Human lung adenocarcinoma cell lines (DeNicola et al., 2015) were maintained in RPMI 1640 medium (Hyclone or Gibco) supplemented with 10% FBS and in the absence of antibiotics at 37°C in a humidified incubator containing 95% air and 5% CO_2_. Cell lines were routinely tested for mycoplasma contamination with the MycoAlert Assay (Lonza). ISCU (addgene #102972), NFS1 (addgene #102963, 102964), and NNT (Open Biosystems TRCN0000028507, TRCN0000028512) knockdown was achieved using validated short hairpin sequences targeting these transcripts in a pLKO.1 vector. Cells were infected with shRNA or scramble lentivirus and then selected with 1 μg/mL puromycin for 3 days prior to each experiment. All experiments were conducted 4-days post lentiviral infection except for analyses of proliferation or viability. The Pos5p nucleotide sequence was purchased as a gBlock from Integrated DNA Technologies®. Pos5p was modified to include flanking 3X HA-tags and cloned into the pLenti-CMV-Blast vector (addgene #17445). Cells were infected with either pos5p or GFP control lentivirus and then selected with 10 μg/mL blasticidin for 5 days. Similar to the strategy used to generate a mitochondrially-targeted catalase expressing mouse (Schriner et al., 2005), the initiating methionine codon of human catalase cDNA was replaced with the sequence for the first 25 amino acids of the ornithine transcarbamylase leader sequence to target exogenous catalase protein to the mitochondria. The MitoCatalase sequence was cloned into the pPHAGE C-TAP vector (Huttlin et al., 2015). Cells were infected with either MitoCatalase or luciferase (Luc) control lentivirus and then selected with 1 μg/mL puromycin for 5 days.

### Analyses of Cell Proliferation and Viability

For proliferation assays, cells were seeded in triplicate on four 96-well plates at a density of 2,500-5,000 cells/well in 100μL. The next day, cells were infected with scramble or shNNT lentivirus in a final volume of 50μL, then overlaid with 100μL of medium and allowed to proliferate. On day 1 post-infection, one of the plates was collected and the cells fixed with 4% paraformaldehyde. This was repeated on days 3, 5, and 7 post-infection. Fixed cells were then stained with crystal violet, washed, and dried overnight. Crystal violet was solubilized with 100μL of 10% acetic acid and the OD_600_ was measured with a spectrophotometer (Promega). For viability assays, cells were plated on 96-well plates at a density of 10,000 cells/well in 100μL. On the next day, cells were incubated with 100μL of media containing tert-butyl hydroperoxide (tbHP; Sigma-Aldrich), cumene hydroperoxide (CHP; Invitrogen), menadione (Fisher Scientific), oleate (Fisher Scientific), or palmitate (Sigma-Aldrich) at the indicated concentrations for 24 hours. Cells were then fixed with 4% paraformaldehyde, stained with crystal violet, washed, and dried overnight. Crystal violet was solubilized with 100μL of 10% acetic acid and the OD_600_ determined. For experiments evaluating the effect of NNT knockdown, cells were seeded on day 2 post-infection. Relative viability was determined following normalization to scramble or DMSO treated cells.

### Immunoblotting

Cell lysates were generated in ice cold RIPA buffer (20mM Tris-HCl [pH 7.5; VWR], 150mM NaCl [Fisher Scientific], 1mM EGTA [VWR], 1mM EDTA [Sigma-Aldrich], 1% sodium deoxycholate [Sigma-Aldrich], 1% NP-40 [Sigma-Aldrich]) supplemented with protease inhibitors (Roche). Protein concentrations were determined by DC Protein Assay (Bio-Rad) prior to mixing with a 6X reducing sample buffer containing β-mercaptoethanol (VWR). Proteins were separated by SDS-PAGE using NuPAGE 4-12% Bis-Tris precast gels (Invitrogen), then transferred to 0.45μM nitrocellulose membranes (GE Healthcare). For analysis of the Prdx3 oxidation state, a previously published protocol was followed (Cox et al., 2009). These redox western samples were mixed with a 4X non-reducing buffer prior to separation by SDS-Page. Membranes were blocked with 5% non-fat milk in TBST and then incubated with the following primary antibodies: ACO2 (GeneTex, GTX109736), Actin (ThermoFisher, AC-15), Catalase (Cell Signaling Technologies, D4P7B), DLST (Cell Signaling Technologies, D22B1), HA-Tag (Cell Signaling Technologies, C29F4), HSP90 (Cell Signaling Technologies, 4874S), ISCU (Santa Cruz Biotechnology, sc-373694), Lipoic acid (Millipore Sigma, 437695), NFS1 (Santa Cruz Biotechnology, sc-365308), NNT for cell lysates (Abcam, ab110352), NNT for mouse tissue lysates (GeneTex, GTX103015), PDH-E2 (Abcam, ab126224), Prdx3 (Abcam, ab73349), α-Tubulin (Santa Crux Biotechnology, sc-8035). HRP-conjugated secondary antibodies and enhanced chemiluminescence were used for all immunoblotting.

### Flow Cytometry Analyses of ROS

For all analyses of ROS, 10^5^ cells were seeded overnight in triplicate wells of a 6-well plate. Mitochondrial H_2_O_2_ levels were determined using the fluorescent dye, MitoPy1 (Tocris), according to an established protocol (Dickinson et al., 2013). Briefly, cells were incubated in fresh media for 4 hours, washed with PBS (Hyclone or Sigma-Aldrich), and incubated in 1mL of 10μM MitoPY1 for 30 minutes. Depending on the experiment, cells were either collected immediately for analysis or challenged with H_2_O_2_ for an additional 30 minutes prior to collection.

Mitochondrial •O_2_^−^ levels were determined with the fluorescent dye, MitoSox Red (Invitrogen), according to an established protocol (Kauffman et al., 2016). Briefly, cells were incubated in fresh media for 4 hours, washed with PBS, and incubated in 1mL of 5μM MitoSOX Red in HBSS (Gibco) for 20 minutes. Cells were then collected immediately for analysis. Cytosolic ROS levels were determined with the fluorescent dye, CellRox Green (Invitrogen), according to the manufacturer’s protocol. Briefly, cells were incubated in 1mL of fresh media for 4 hours, at which point 2μL of 2.5mM CellROX Green was added to each well for a final concentration of 5μM. Cells were incubated with CellROX green for 30 minutes and then collected for analysis. The fluorescence of dye-loaded cells was determined by flow cytometry with a BD Biosciences 2 Laser 4 Color FacsCalibur Flow Cytometer (Marshall Scientific). The FL1 channel was used for analyses of MitoPY1 and CellROX Green fluorescence, whereas the FL3 channel was used for analyses of MitoSOX Red fluorescence. The mean fluorescence intensity of 10,000 discrete events were calculated for each sample.

### NADPH:NADP^+^ Assay

25,000 cells were seeded in 500μL of medium overnight in triplicate on 12-well plates. Cells were then incubated in fresh media for 4 hours, collected, and extracted to determine the NADPH/NADP+ ratio according to the NADP/NADPH-Glo Assay Kit (Promega) protocol.

### Seahorse Analyses of Mitochondrial Function

All measures of oxygen consumption were determined with a Seahorse XFe96 Analyzer (Agilent). General mitochondrial function was assessed according to the Seahorse XF Cell Mito Stress Kit protocol (Agilent). Assessments of individual respiratory chain activities were performed according to a previously established protocol (Salabei et al., 2014). Briefly, 40,000 cells were plated in quadruplicate on an XFe96 microplate and allowed to seed overnight. Immediately prior to assay, cells were overlaid with 175μL of mitochondrial assay solution (220mM mannitol [Sigma-Aldrich], 70mM sucrose [Sigma-Aldrich], 10mM KH_2_PO_4_ [VWR], 5mM MgCl_2_ [VWR], 2mM HEPES [Fisher Scientific], 1mM EGTA [VWR]) supplemented with the Seahorse Plasma Membrane Permeabilizer (Agilent), 4mM ADP (Sigma-Aldrich), and either 10mM sodium pyruvate (Sigma-Aldrich) and 1mM malate (Sigma-Aldrich) or 10mM glutamate (Sigma-Aldrich) and 1mM malate. Cells were then sequentially subjected to 2μM rotenone (Sigma-Aldrich), 10mM succinate (Sigma-Aldrich), 2μM antimycin A (Sigma-Aldrich), and 10mM ascorbate (Sigma-Aldrich) with 100μM N,N,N’,N’-tetramethyl-ρ-phenylene diamine (Sigma-Aldrich). Lastly, palmitate oxidation was determined using the XF Fatty Acid Oxidation Assay Kit (Agilent) protocol.

### Aconitase Assay

Aconitase activity was determined based on a modified version of a protocol previously described (Francisco et al., 2018). Cells were collected and resuspended in 250μL of 50mM Tris-HCl, 150mM NaCl, pH 7.4. The cell suspension was homogenized with a dounce homogenizer and the homogenate spun down for 10 minutes at 10,000xg at 4°C. The pellet was then washed twice and resuspended in 100μL of 1% triton (Sigma-Aldrich) in 50mM Tris-HCl, pH 7.4 to lyse the mitochondrial membrane. This fraction was then spun down for 15 minutes at 17,000xg at 4°C. The protein concentration was then determined by DC Protein Assay (Bio-Rad) and 175μL of 100-500ug/mL protein solution was generated with assay buffer (50mM Tris-HCl, pH 7.4). 50μL of this solution was transferred to triplicate wells of a black-walled 96-well fluorescence microplate already containing 55μL of assay buffer. Next, 50μL of a 4mM NADP^+^ (Sigma-Aldrich), 20U/mL IDH1 (Sigma-Aldrich) solution was added to each well. Finally, 50μL of 10mM sodium citrate (Sigma-Aldrich) was added to each well to initiate the assay. The plate was transferred to a fluorescence-compatible plate reader (Promega) to measure NADPH autofluorescence every minute over a period of an hour. This change in fluorescence over time is indicative of aconitase activity, where ACO2 present in the mitochondrial protein fraction converts the supplied citrate to isocitrate, which the supplied IDH1 then metabolizes in a reaction that generates NADPH. For the analysis of tumor tissue, tumors were homogenized in 500μL of mitochondrial isolation buffer (200mM mannitol, 10mM sucrose, 1mM EGTA, 10mM HEPES, pH 7.4). Homogenates were spun down for 10 minutes at 800xg at 4°C. The supernatant was subjected to an additional spin of 10,000xg at 4°C. The pellet was then processed for analysis as described.

### LC-HRMS Metabolomics

NSCLC cells seeded in triplicate wells of a 6-well plate were rapidly washed in ice cold PBS and extracted in 0.5mL of 80% methanol overnight at −80°C. Extracts were then cleared by centrifugation (17,000xg for 30 minutes at 4°C), and the supernatant analyzed by liquid chromatography-high resolution mass spectrometry (LC-HRMS). We performed this LC-HRMS analysis under the conditions for non-targeted metabolomics that we have established previously (Kang et al., 2019). Briefly, we utilized a Vanquish UPLC system coupled to a Q Exactive HF mass spectrometer equipped with heated electrospray ionization (Thermo Fisher Scientific). A SeQuant ZIC-pHILIC LC column, 5μm, 150 × 4.6mm (Millipore Sigma) with a SeQuant ZIC-pHILIC guard column, 20 × 4.6mm (Millipore Sigma) was used for chromatographic separation. The mobile phase A consisted of 10mM ammonium carbonate and 0.05% ammonium hydroxide in water, while mobile phase B was 100% acetonitrile. The MS1 scan was operated in both positive and negative mode for data acquisition and data was analyzed with El Maven v0.3.1 (Clasquin et al., 2012). Metabolite identification was based on a comparison of both the m/z value and retention time of sample peaks to an internal MSMLS library (Sigma-Aldrich).

### Statistical Analysis

Data were analyzed for statistical significance with GraphPad Prism 8 software. Values of p<0.05 were considered significant (n.s., not significant; *p<0.05; **p<0.01; ***p<0.001; ****p<0.0001). Differences between survival curves were determined by the Log-rank test. Comparisons of two groups were performed with a two-sided unpaired Student’s t test. A one-way ANOVA with a post-hoc Brown-Forsythe test was performed for comparisons of 3 or more groups, as similar variances between groups were observed. Data reported as mean ± SD of at least three technical replicates and representative of at least three experimental replicates unless noted otherwise. In assessing the effects of NNT knockdown in GFP/pos5p and Luc/MitoCatalase cells a two-way ANOVA with a Sidak’s multiple comparisons test was performed.

## Supplemental material

Fig. S1 shows analyses of ROS levels and the response to exogenous oxidant treatment in NSCLC cells subject to NNT knockdown. Fig. S2 shows additional measures of mitochondrial function in NNT-deficient NSCLC cells in addition to immunoblot analyses of ACO2 expression in NNT-deficient NSCLC cells and KP tumors with differential NNT expression. Fig. S3 shows analyses of basal mitochondrial function in pos5p expressing NSCLC cells. Fig. S4 demonstrates the functionality of MitoCatalase in MitoCatalase expressing NSCLC cells.

## Author contributions

N.P. Ward designed and performed the experiments, analyzed the data, and wrote the manuscript; Y.P. Kang performed the LC-HRMS analysis; A. Falzone maintained mice cohorts, performed adenoviral-Cre infections, and collected lung tissue; T.A. Boyle provided pathology expertise for evaluation of the lung sections; G.M. DeNicola conceived of the study, contributed to the experimental design, performed adenoviral-Cre infections, collected lung tissue, and wrote the manuscript. All authors reviewed the data and commented on the manuscript.

## Acknowledgments

We would like to thank Janine DeBlasi and Sarah Naomi Olsen for technical assistance and members of the DeNicola lab for their helpful discussion. This work was supported by grants from the V Foundation (V2017-015), NIH (R37-CA230042) and Hope Funds for Cancer Research to G.M.D, and the AACR-Takeda Oncology Lung Cancer Research Fellowship (19-40-38-KANG) to Y.P.K. This work was also supported by the Flow Cytometry Core and the Proteomics/Metabolomics care, which are funded in part by Moffitt’s Cancer Center Support Grant (NCI, P30-CA076292), and grants from the Moffitt Foundation, and a Florida Bankhead-Coley grant (06BS-02–9614) to the Proteomics/Metabolomics core.

The authors declare no competing financial interests.

## Abbreviations

•O_2_^−^: Superoxide
ACO2: Aconitase 2
CHP: Cumene Hydroperoxide
DLST: Dihydrolipoamide S-Succinyltransferase
ETC: Electron Transport Chain
Fe-S: Iron-Sulfur
GEMM: Genetically Engineered Mouse Model
H_2_O_2_: Hydrogen Peroxide
IMM: Inner Mitochondrial Membrane
ISCU: Iron Sulfur Scaffold Protein
KP: LSL-Kras^G12D^; Trp53^flox/flox^
NADH: Nicotinamide Adenine Dinucleotide
NADPH: Nicotinamide Adenine Dinucleotide Phosphate
NFS1: Cysteine Desulfurase
NNT: Nicotinamide Nucleotide Transhydrogenase
NSCLC: Non-Small Cell Lung Cancer
OCR: Oxygen Consumption Rate
PDH: Pyruvate Dehydrogenase
tbHP: Tert-butyl Hydroperoxide

## References

Alhebshi, A., Sideri, T. C., Holland, S. L. & Avery, S. V. 2012. The essential iron-sulfur protein Rli1 is an important target accounting for inhibition of cell growth by reactive oxygen species. Mol Biol Cell, 23, 3582–90.

Alvarez, S. W., Sviderskiy, V. O., Terzi, E. M., Papagiannakopoulos, T., Moreira, A. L., Adams, S., Sabatini, D. M., Birsoy, K. & Possemato, R. 2017. NFS1 undergoes positive selection in lung tumours and protects cells from ferroptosis. Nature, 551, 639–643.

Chortis, V., Taylor, A. E., Doig, C. L., Walsh, M. D., Meimaridou, E., Jenkinson, C., Rodriguez-Blanco, G., Ronchi, C. L., Jafri, A., Metherell, L. A., Hebenstreit, D., Dunn, W. B., Arlt, W. & Foster, P. A. 2018. Nicotinamide Nucleotide Transhydrogenase as a Novel Treatment Target in Adrenocortical Carcinoma. Endocrinology, 159, 2836–2849.

Clasquin, M. F., Melamud, E. & Rabinowitz, J. D. 2012. LC-MS data processing with MAVEN: a metabolomic analysis and visualization engine. Curr Protoc Bioinformatics, Chapter 14, Unit14 11.

Cox, A. G., Pearson, A. G., Pullar, J. M., Jonsson, T. J., Lowther, W. T., Winterbourn, C. C. & Hampton, M. B. 2009. Mitochondrial peroxiredoxin 3 is more resilient to hyperoxidation than cytoplasmic peroxiredoxins. Biochem J, 421, 51–8.

Crack, J. C., Green, J., Thomson, A. J. & Le Brun, N. E. 2014. Iron-sulfur clusters as biological sensors: the chemistry of reactions with molecular oxygen and nitric oxide. Acc Chem Res, 47, 3196–205.

Crooks, D. R., Maio, N., Lane, A. N., Jarnik, M., Higashi, R. M., Haller, R. G., Yang, Y., Fan, T. W., Linehan, W. M. & Rouault, T. A. 2018. Acute loss of iron-sulfur clusters results in metabolic reprogramming and generation of lipid droplets in mammalian cells. J Biol Chem, 293, 8297–8311.

D’souza, S. F. & Srere, P. A. 1983. Binding of citrate synthase to mitochondrial inner membranes. J Biol Chem, 258, 4706–9.

Davidson, S. M., Papagiannakopoulos, T., Olenchock, B. A., Heyman, J. E., Keibler, M. A., Luengo, A., Bauer, M. R., Jha, A. K., O’brien, J. P., Pierce, K. A., Gui, D. Y., Sullivan, L. B., Wasylenko, T. M., Subbaraj, L., Chin, C. R., Stephanopolous, G., Mott, B. T., Jacks, T., Clish, C. B. & Vander Heiden, M. G. 2016. Environment Impacts the Metabolic Dependencies of Ras-Driven Non-Small Cell Lung Cancer. Cell Metab, 23, 517–28.

Denicola, G. M., Chen, P. H., Mullarky, E., Sudderth, J. A., Hu, Z., Wu, D., Tang, H., Xie, Y., Asara, J. M., Huffman, K. E., Wistuba, II, Minna, J. D., Deberardinis, R. J. & Cantley, L. C. 2015. NRF2 regulates serine biosynthesis in non-small cell lung cancer. Nat Genet, 47, 1475–81.

Denicola, G. M., Karreth, F. A., Humpton, T. J., Gopinathan, A., Wei, C., Frese, K., Mangal, D., Yu, K. H., Yeo, C. J., Calhoun, E. S., Scrimieri, F., Winter, J. M., Hruban, R. H., Iacobuzio-Donahue, C., Kern, S. E., Blair, I. A. & Tuveson, D. A. 2011. Oncogene-induced Nrf2 transcription promotes ROS detoxification and tumorigenesis. Nature, 475, 106–9.

Dickinson, B. C., Lin, V. S. & Chang, C. J. 2013. Preparation and use of MitoPY1 for imaging hydrogen peroxide in mitochondria of live cells. Nat Protoc, 8, 1249–59.

Djaman, O., Outten, F. W. & Imlay, J. A. 2004. Repair of oxidized iron-sulfur clusters in Escherichia coli. J Biol Chem, 279, 44590–9.

Ducker, G. S., Chen, L., Morscher, R. J., Ghergurovich, J. M., Esposito, M., Teng, X., Kang, Y. & Rabinowitz, J. D. 2016. Reversal of Cytosolic One-Carbon Flux Compensates for Loss of the Mitochondrial Folate Pathway. Cell Metab, 23, 1140–1153.

Dupage, M., Dooley, A. L. & Jacks, T. 2009. Conditional mouse lung cancer models using adenoviral or lentiviral delivery of Cre recombinase. Nat Protoc, 4, 1064–72.

Fan, T. W., Lane, A. N., Higashi, R. M., Farag, M. A., Gao, H., Bousamra, M. & Miller, D. M. 2009. Altered regulation of metabolic pathways in human lung cancer discerned by (13)C stable isotope-resolved metabolomics (SIRM). Mol Cancer, 8, 41.

Faubert, B., Li, K. Y., Cai, L., Hensley, C. T., Kim, J., Zacharias, L. G., Yang, C., Do, Q. N., Doucette, S., Burguete, D., Li, H., Huet, G., Yuan, Q., Wigal, T., Butt, Y., Ni, M., Torrealba, J., Oliver, D., Lenkinski, R. E., Malloy, C. R., Wachsmann, J. W., Young, J. D., Kernstine, K. & Deberardinis, R. J. 2017. Lactate Metabolism in Human Lung Tumors. Cell, 171, 358–371 e9.

Fernie, A. R., Zhang, Y. & Sweetlove, L. J. 2018. Passing the Baton: Substrate Channelling in Respiratory Metabolism. Research, 2018, 16.

Fisher-Wellman, K. H., Gilliam, L. A. A., Lin, C. T., Cathey, B. L., Lark, D. S. & Darrell Neufer, P. 2013. Mitochondrial glutathione depletion reveals a novel role for the pyruvate dehydrogenase complex as a key H2O2-emitting source under conditions of nutrient overload. Free Radic Biol Med, 65, 1201–1208.

Fisher-Wellman, K. H., Lin, C. T., Ryan, T. E., Reese, L. R., Gilliam, L. A., Cathey, B. L., Lark, D. S., Smith, C. D., Muoio, D. M. & Neufer, P. D. 2015. Pyruvate dehydrogenase complex and nicotinamide nucleotide transhydrogenase constitute an energy-consuming redox circuit. Biochem J, 467, 271–80.

Flint, D. H., Tuminello, J. F. & Emptage, M. H. 1993. The inactivation of Fe-S cluster containing hydro-lyases by superoxide. J Biol Chem, 268, 22369–76.

Fosset, C., Chauveau, M. J., Guillon, B., Canal, F., Drapier, J. C. & Bouton, C. 2006. RNA silencing of mitochondrial m-Nfs1 reduces Fe-S enzyme activity both in mitochondria and cytosol of mammalian cells. J Biol Chem, 281, 25398–406.

Francisco, A., Ronchi, J. A., Navarro, C. D. C., Figueira, T. R. & Castilho, R. F. 2018. Nicotinamide nucleotide transhydrogenase is required for brain mitochondrial redox balance under hampered energy substrate metabolism and high-fat diet. J Neurochem, 147, 663–677.

Gameiro, P. A., Laviolette, L. A., Kelleher, J. K., Iliopoulos, O. & Stephanopoulos, G. 2013. Cofactor balance by nicotinamide nucleotide transhydrogenase (NNT) coordinates reductive carboxylation and glucose catabolism in the tricarboxylic acid (TCA) cycle. J Biol Chem, 288, 12967–77.

Hensley, C. T., Faubert, B., Yuan, Q., Lev-Cohain, N., Jin, E., Kim, J., Jiang, L., Ko, B., Skelton, R., Loudat, L., Wodzak, M., Klimko, C., Mcmillan, E., Butt, Y., Ni, M., Oliver, D., Torrealba, J., Malloy, C. R., Kernstine, K., Lenkinski, R. E. & Deberardinis, R. J. 2016. Metabolic Heterogeneity in Human Lung Tumors. Cell, 164, 681–94.

Ho, H. Y., Lin, Y. T., Lin, G., Wu, P. R. & Cheng, M. L. 2017. Nicotinamide nucleotide transhydrogenase (NNT) deficiency dysregulates mitochondrial retrograde signaling and impedes proliferation. Redox Biol, 12, 916–928.

Huttlin, E. L., Ting, L., Bruckner, R. J., Gebreab, F., Gygi, M. P., Szpyt, J., Tam, S., Zarraga, G., Colby, G., Baltier, K., Dong, R., Guarani, V., Vaites, L. P., Ordureau, A., Rad, R., Erickson, B. K., Wuhr, M., Chick, J., Zhai, B., Kolippakkam, D., Mintseris, J., Obar, R. A., Harris, T., Artavanis-Tsakonas, S., Sowa, M. E., De CAMILLI, P., Paulo, J. A., Harper, J. W. & Gygi, S. P. 2015. The BioPlex Network: A Systematic Exploration of the Human Interactome. Cell, 162, 425–440.

Jackson, E. L., Olive, K. P., Tuveson, D. A., Bronson, R., Crowley, D., Brown, M. & Jacks, T. 2005. The differential effects of mutant p53 alleles on advanced murine lung cancer. Cancer Res, 65, 10280–8.

Jackson, E. L., Willis, N., Mercer, K., Bronson, R. T., Crowley, D., Montoya, R., Jacks, T. & Tuveson, D. A. 2001. Analysis of lung tumor initiation and progression using conditional expression of oncogenic K-ras. Genes Dev, 15, 3243–8.

Jiang, L., Shestov, A. A., Swain, P., Yang, C., Parker, S. J., Wang, Q. A., Terada, L. S., Adams, N. D., Mccabe, M. T., Pietrak, B., Schmidt, S., Metallo, C. M., Dranka, B. P., Schwartz, B. & Deberardinis, R. J. 2016. Reductive carboxylation supports redox homeostasis during anchorage-independent growth. Nature, 532, 255–8.

Johnson, D. C., Dean, D. R., Smith, A. D. & Johnson, M. K. 2005. Structure, function, and formation of biological iron-sulfur clusters. Annu Rev Biochem, 74, 247–81.

Kampjut, D. & Sazanov, L. A. 2019. Structure and mechanism of mitochondrial proton-translocating transhydrogenase. Nature.

Kang, Y. P., Torrente, L., Falzone, A., Elkins, C. M., Liu, M., Asara, J. M., Dibble, C. C. & Denicola, G. M. 2019. Cysteine dioxygenase 1 is a metabolic liability for non-small cell lung cancer. Elife, 8.

Kauffman, M. E., Kauffman, M. K., Traore, K., Zhu, H., Trush, M. A., Jia, Z. & Li, Y. R. 2016. MitoSOX-Based Flow Cytometry for Detecting Mitochondrial ROS. React Oxyg Species (Apex), 2, 361–370.

Klingenberg, M. & Slenczka, W. 1959. [Pyridine nucleotide in liver mitochondria. An analysis of their redox relationships]. Biochem Z, 331, 486–517.

Law, L. W., Ting, R. C. & Leckband, E. 1967. Prevention of virus-induced neoplasms in mice through passive transfer of immunity by sensitized syngeneic lymphoid cells. Proc Natl Acad Sci U S A, 57, 1068–75.

Lee, W. H., Kim, J. W., Park, E. H., Han, N. S., Kim, M. D. & Seo, J. H. 2013. Effects of NADH kinase on NADPH-dependent biotransformation processes in Escherichia coli. Appl Microbiol Biotechnol, 97, 1561–9.

Li, S., Zhuang, Z., Wu, T., Lin, J. C., Liu, Z. X., Zhou, L. F., Dai, T., Lu, L. & Ju, H. Q. 2018. Nicotinamide nucleotide transhydrogenase-mediated redox homeostasis promotes tumor growth and metastasis in gastric cancer. Redox Biol, 18, 246–255.

Lill, R. & Muhlenhoff, U. 2008. Maturation of iron-sulfur proteins in eukaryotes: mechanisms, connected processes, and diseases. Annu Rev Biochem, 77, 669–700.

Lopert, P. & Patel, M. 2014. Nicotinamide nucleotide transhydrogenase (Nnt) links the substrate requirement in brain mitochondria for hydrogen peroxide removal to the thioredoxin/peroxiredoxin (Trx/Prx) system. J Biol Chem, 289, 15611–20.

Mayers, J. R., Torrence, M. E., Danai, L. V., Papagiannakopoulos, T., Davidson, S. M., Bauer, M. R., Lau, A. N., Ji, B. W., Dixit, P. D., Hosios, A. M., Muir, A., Chin, C. R., Freinkman, E., Jacks, T., Wolpin, B. M., Vitkup, D. & Vander HEIDEN, M. G. 2016. Tissue of origin dictates branched-chain amino acid metabolism in mutant Kras-driven cancers. Science, 353, 1161–5.

Meimaridou, E., Goldsworthy, M., Chortis, V., Fragouli, E., Foster, P. A., Arlt, W., Cox, R. & Metherell, L. A. 2018. NNT is a key regulator of adrenal redox homeostasis and steroidogenesis in male mice. J Endocrinol, 236, 13–28.

Meuwissen, R. & Berns, A. 2005. Mouse models for human lung cancer. Genes Dev, 19, 643–64.

Migita, T., Narita, T., Nomura, K., Miyagi, E., Inazuka, F., Matsuura, M., Ushijima, M., Mashima, T., Seimiya, H., Satoh, Y., Okumura, S., Nakagawa, K. & Ishikawa, Y. 2008. ATP citrate lyase: activation and therapeutic implications in non-small cell lung cancer. Cancer Res, 68, 8547–54.

Miller, J. R., Busby, R. W., Jordan, S. W., Cheek, J., Henshaw, T. F., Ashley, G. W., Broderick, J. B., Cronan, J. E., JR. & Marletta, M. A. 2000. Escherichia coli LipA is a lipoyl synthase: in vitro biosynthesis of lipoylated pyruvate dehydrogenase complex from octanoyl-acyl carrier protein. Biochemistry, 39, 15166–78.

Morgunov, I. & Srere, P. A. 1998. Interaction between citrate synthase and malate dehydrogenase. Substrate channeling of oxaloacetate. J Biol Chem, 273, 29540–4.

Mullen, A. R., Hu, Z., Shi, X., Jiang, L., Boroughs, L. K., Kovacs, Z., Boriack, R., Rakheja, D., Sullivan, L. B., Linehan, W. M., Chandel, N. S. & Deberardinis, R. J. 2014. Oxidation of alpha-ketoglutarate is required for reductive carboxylation in cancer cells with mitochondrial defects. Cell Rep, 7, 1679–1690.

Navarro, C. D. C., Figueira, T. R., Francisco, A., Dal’BO, G. A., Ronchi, J. A., Rovani, J. C., Escanhoela, C. A. F., Oliveira, H. C. F., Castilho, R. F. & Vercesi, A. E. 2017. Redox imbalance due to the loss of mitochondrial NAD(P)-transhydrogenase markedly aggravates high fat diet-induced fatty liver disease in mice. Free Radic Biol Med, 113, 190–202.

Padanad, M. S., Konstantinidou, G., Venkateswaran, N., Melegari, M., Rindhe, S., Mitsche, M., Yang, C., Batten, K., Huffman, K. E., Liu, J., Tang, X., Rodriguez-Canales, J., Kalhor, N., Shay, J. W., Minna, J. D., Mcdonald, J., Wistuba, II, Deberardinis, R. J. & Scaglioni, P. P. 2016. Fatty Acid Oxidation Mediated by Acyl-CoA Synthetase Long Chain 3 Is Required for Mutant KRAS Lung Tumorigenesis. Cell Rep, 16, 1614–1628.

Pain, J., Balamurali, M. M., Dancis, A. & Pain, D. 2010. Mitochondrial NADH kinase, Pos5p, is required for efficient iron-sulfur cluster biogenesis in Saccharomyces cerevisiae. J Biol Chem, 285, 39409–24.

Parry, R. J. & Trainor, D. A. 1978. Biosynthesis of lipoic acid. 2. Stereochemistry of sulfur introduction at C-6 of octanoic acid. Journal of the American Chemical Society, 100, 5243–5244.

Porpaczy, Z., Sumegi, B. & Alkonyi, I. 1987. Interaction between NAD-dependent isocitrate dehydrogenase, alpha-ketoglutarate dehydrogenase complex, and NADH:ubiquinone oxidoreductase. J Biol Chem, 262, 9509–14.

Porporato, P. E., Filigheddu, N., Pedro, J. M. B., Kroemer, G. & Galluzzi, L. 2018. Mitochondrial metabolism and cancer. Cell Res, 28, 265–280.

Rao, S., Mondragon, L., Pranjic, B., Hanada, T., Stoll, G., Kocher, T., Zhang, P., Jais, A., Lercher, A., Bergthaler, A., Schramek, D., Haigh, K., Sica, V., Leduc, M., Modjtahedi, N., Pai, T. P., Onji, M., Uribesalgo, I., Hanada, R., Kozieradzki, I., Koglgruber, R., Cronin, S. J., She, Z., Quehenberger, F., Popper, H., Kenner, L., Haigh, J. J., Kepp, O., Rak, M., Cai, K., Kroemer, G. & Penninger, J. M. 2019. AIF-regulated oxidative phosphorylation supports lung cancer development. Cell Res, 29, 579–591.

Ren, J. G., Seth, P., Clish, C. B., Lorkiewicz, P. K., Higashi, R. M., Lane, A. N., Fan, T. W. & Sukhatme, V. P. 2014. Knockdown of malic enzyme 2 suppresses lung tumor growth, induces differentiation and impacts PI3K/AKT signaling. Sci Rep, 4, 5414.

Romero, R., Sayin, V. I., Davidson, S. M., Bauer, M. R., Singh, S. X., Leboeuf, S. E., Karakousi, T. R., Ellis, D. C., Bhutkar, A., Sanchez-Rivera, F. J., Subbaraj, L., Martinez, B., Bronson, R. T., Prigge, J. R., Schmidt, E. E., Thomas, C. J., Goparaju, C., Davies, A., Dolgalev, I., Heguy, A., Allaj, V., Poirier, J. T., Moreira, A. L., Rudin, C. M., Pass, H. I., Vander HEIDEN, M. G., Jacks, T. & Papagiannakopoulos, T. 2017. Keap1 loss promotes Kras-driven lung cancer and results in dependence on glutaminolysis. Nat Med, 23, 1362–1368.

Ronchi, J. A., Francisco, A., Passos, L. A., Figueira, T. R. & Castilho, R. F. 2016. The Contribution of Nicotinamide Nucleotide Transhydrogenase to Peroxide Detoxification Is Dependent on the Respiratory State and Counterbalanced by Other Sources of NADPH in Liver Mitochondria. J Biol Chem, 291, 20173–87.

Rouault, T. A. 2015. Mammalian iron-sulphur proteins: novel insights into biogenesis and function. Nat Rev Mol Cell Biol, 16, 45–55.

Roucher-Boulez, F., Mallet-Motak, D., Samara-Boustani, D., Jilani, H., Ladjouze, A., Souchon, P. F., Simon, D., Nivot, S., Heinrichs, C., Ronze, M., Bertagna, X., Groisne, L., Leheup, B., Naud-Saudreau, C., Blondin, G., Lefevre, C., Lemarchand, L. & Morel, Y. 2016. NNT mutations: a cause of primary adrenal insufficiency, oxidative stress and extra-adrenal defects. Eur J Endocrinol, 175, 73–84.

Rydstrom, J. 2006. Mitochondrial NADPH, transhydrogenase and disease. Biochim Biophys Acta, 1757, 721–6.

Salabei, J. K., Gibb, A. A. & Hill, B. G. 2014. Comprehensive measurement of respiratory activity in permeabilized cells using extracellular flux analysis. Nat Protoc, 9, 421–38.

Sauer, U., Canonaco, F., Heri, S., Perrenoud, A. & Fischer, E. 2004. The soluble and membrane-bound transhydrogenases UdhA and PntAB have divergent functions in NADPH metabolism of Escherichia coli. J Biol Chem, 279, 6613–9.

Schriner, S. E., Linford, N. J., Martin, G. M., Treuting, P., Ogburn, C. E., Emond, M., Coskun, P. E., Ladiges, W., Wolf, N., Van REMMEN, H., Wallace, D. C. & Rabinovitch, P. S. 2005. Extension of murine life span by overexpression of catalase targeted to mitochondria. Science, 308, 1909–11.

Sellers, K., Fox, M. P., Bousamra, M., 2nd, Slone, S. P., Higashi, R. M., Miller, D. M., Wang, Y., Yan, J., Yuneva, M. O., Deshpande, R., Lane, A. N. & Fan, T. W. 2015. Pyruvate carboxylase is critical for non-small-cell lung cancer proliferation. J Clin Invest, 125, 687–98.

Singh, A., Misra, V., Thimmulappa, R. K., Lee, H., Ames, S., Hoque, M. O., Herman, J. G., Baylin, S. B., Sidransky, D., Gabrielson, E., Brock, M. V. & Biswal, S. 2006. Dysfunctional KEAP1-NRF2 interaction in non-small-cell lung cancer. PLoS Med, 3, e420.

Sumegi, B. & Srere, P. A. 1984a. Binding of the enzymes of fatty acid beta-oxidation and some related enzymes to pig heart inner mitochondrial membrane. J Biol Chem, 259, 8748–52.

Sumegi, B. & Srere, P. A. 1984b. Complex I binds several mitochondrial NAD-coupled dehydrogenases. J Biol Chem, 259, 15040–5.

Toye, A. A., Lippiat, J. D., Proks, P., Shimomura, K., Bentley, L., Hugill, A., Mijat, V., Goldsworthy, M., Moir, L., Haynes, A., Quarterman, J., Freeman, H. C., Ashcroft, F. M. & Cox, R. D. 2005. A genetic and physiological study of impaired glucose homeostasis control in C57BL/6J mice. Diabetologia, 48, 675–86.

Webert, H., Freibert, S. A., Gallo, A., Heidenreich, T., Linne, U., Amlacher, S., Hurt, E., Muhlenhoff, U., Banci, L. & Lill, R. 2014. Functional reconstitution of mitochondrial Fe/S cluster synthesis on Isu1 reveals the involvement of ferredoxin. Nat Commun, 5, 5013.

Weinberg, F., Hamanaka, R., Wheaton, W. W., Weinberg, S., Joseph, J., Lopez, M., Kalyanaraman, B., Mutlu, G. M., Budinger, G. R. & Chandel, N. S. 2010. Mitochondrial metabolism and ROS generation are essential for Kras-mediated tumorigenicity. Proc Natl Acad Sci U S A, 107, 8788–93.

Weinberg, S. E. & Chandel, N. S. 2015. Targeting mitochondria metabolism for cancer therapy. Nat Chem Biol, 11, 9–15.

