## Supplemental Figures for "NNT Regulates Mitochondrial Metabolism in NSCLC Through Maintenance of Fe-S Protein Function"

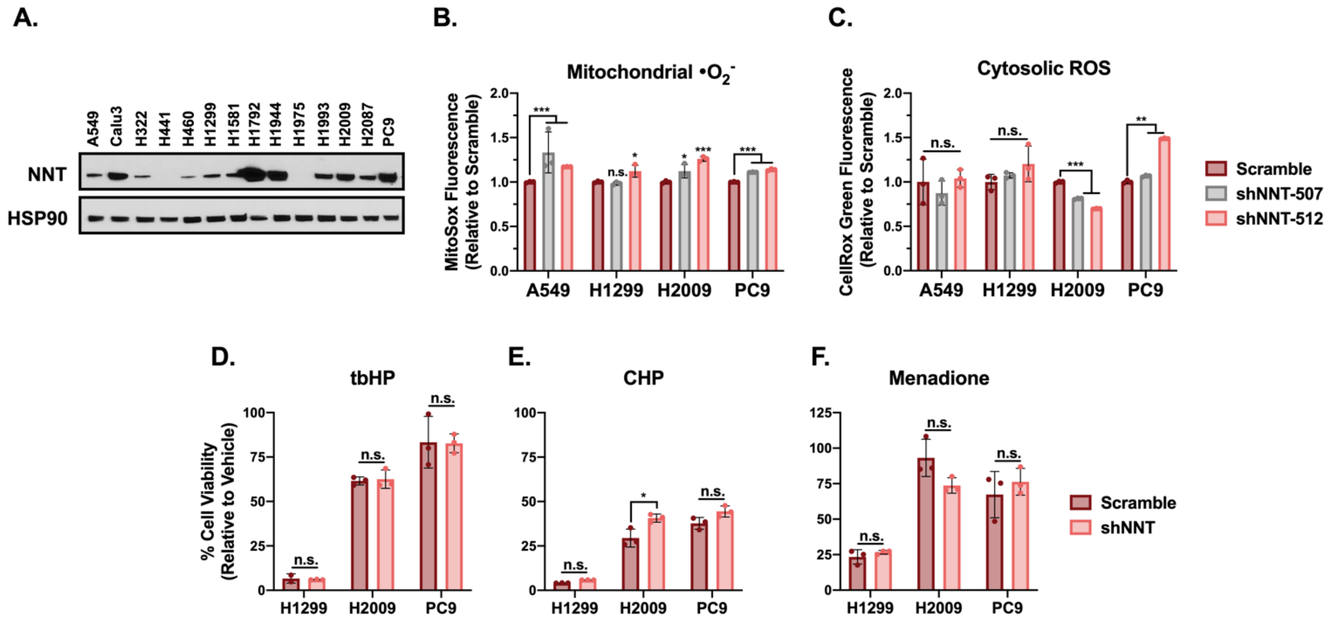


Figure S1. **NNT loss does not sensitize NSCLC cells to exogenous oxidants. (A)** Immunoblot analysis of NNT and HSP90 (loading control) expression in a panel of human NSCLC cells. **(B)** Fluorescence of the mitochondrial $\bullet O_2^-$ sensitive dye, MitoSOX Red, in NSCLC cells following NNT knockdown, relative to scramble infected control cells (one-way ANOVA). **(C)** Fluorescence of the cytosolic ROS sensitive dye, CellROX Green, in NSCLC cells following NNT knockdown, relative to scramble infected control cells (one-way ANOVA). **(D-F)** Viability of NSCLC cells subject to scramble or shNNT lentiviral infection following 24-hour treatment with 15μM (D) tbHP, (E) CHP, or (F) menadione (Student's t test). Cell viability was determined relative to vehicle treated controls. Data are representative of one experiment of three experimental replicates. For B-F, data are represented as mean ± SD of three technical replicates. n.s., not significant; *, p<0.05; **, p<0.01; ***, p<0.001.

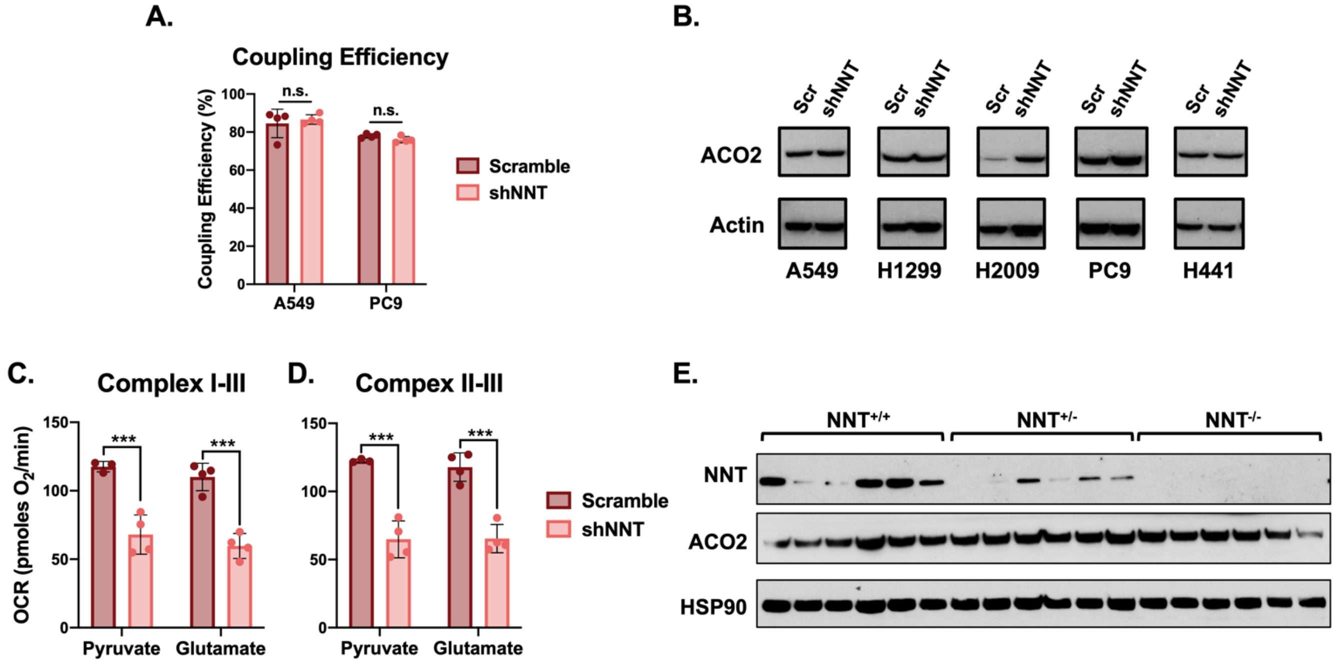


Figure S2. **The effect of NNT loss on ACO2 is independent of changes in protein expression. (A)** Average coupling efficiency of A549 and PC9 cells following infection with either scramble or shNNT lentivirus (Student's t test). **(B)** Immunoblot analysis of ACO2 and actin (loading control) expression in NSCLC cells subject to NNT knockdown. **(C)** Average complex I-III activity following stimulation with 1mM malate and either 10mM pyruvate or 10mM glutamate in H2009 cells subject to NNT knockdown (one-way ANOVA). **(D)** Average complex II-III activity in H2009 cells subject to NNT knockdown following stimulation with 10mM succinate in the presence of 1mM malate and either 10mM pyruvate or 10mM glutamate (one-way ANOVA). **(E)** Immunoblot analysis of NNT, ACO2, and HSP90 (loading control) expression in lung tumors collected from LSL-Kras$^{G12D/+}$; Trp53$^{flox/flox}$; NNT$^{+/+}$ (n=6), LSL-Kras$^{G12D/+}$; Trp53$^{flox/flox}$; NNT$^{+/-}$ (n=6), and LSL-Kras$^{G12D/+}$; Trp53$^{flox/flox}$; NNT$^{-/-}$ (n=6) mice. Data are representative of one experiment of three experimental replicates. For A, C, and D, data are represented as mean $\pm$ SD of at least three technical replicates. n.s., not significant; ***, p<0.001.

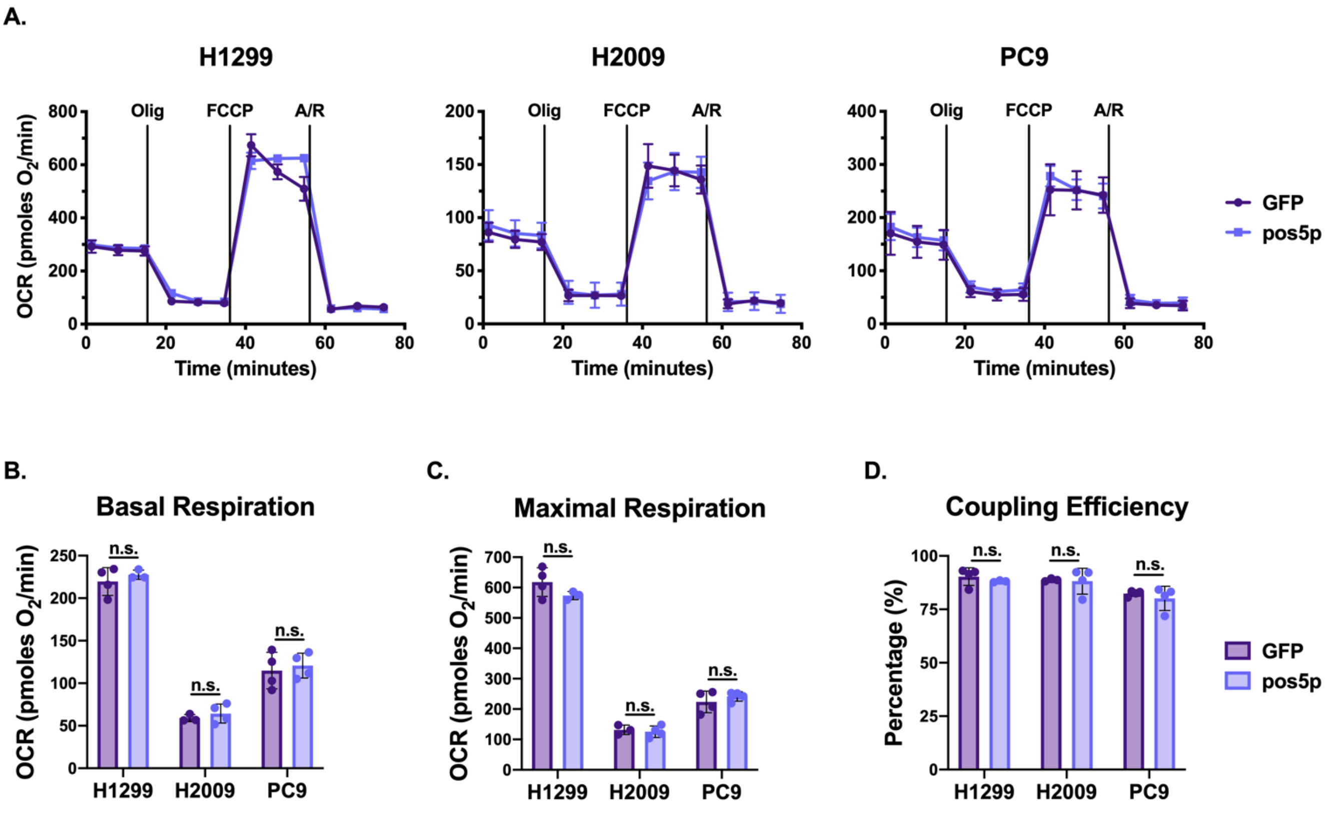


Figure S3. **Pos5p expression does not disrupt mitochondrial function in NSCLC cells. (A)** Plots of OCR in GFP or pos5p expressing NSCLC cells. Cells were supplemented with 10mM glucose and 1mM glutamine and then sequentially challenged with 1μM oligomycin (Olig), 0.5μM of carbonyl cyanide 4-(trifluoromethoxy)phenylhydrazone (FCCP), and 1μM each of antimycin A (A) and rotenone (R). **(B-D)** Average measures of (B) basal respiration, (C) maximal respiration, and (D) coupling efficiency in GFP or pos5p expressing NSCLC cells. Data are representative of one experiment of three experimental replicates. Data are represented as mean ± SD of at least three technical replicates. n.s., not significant.

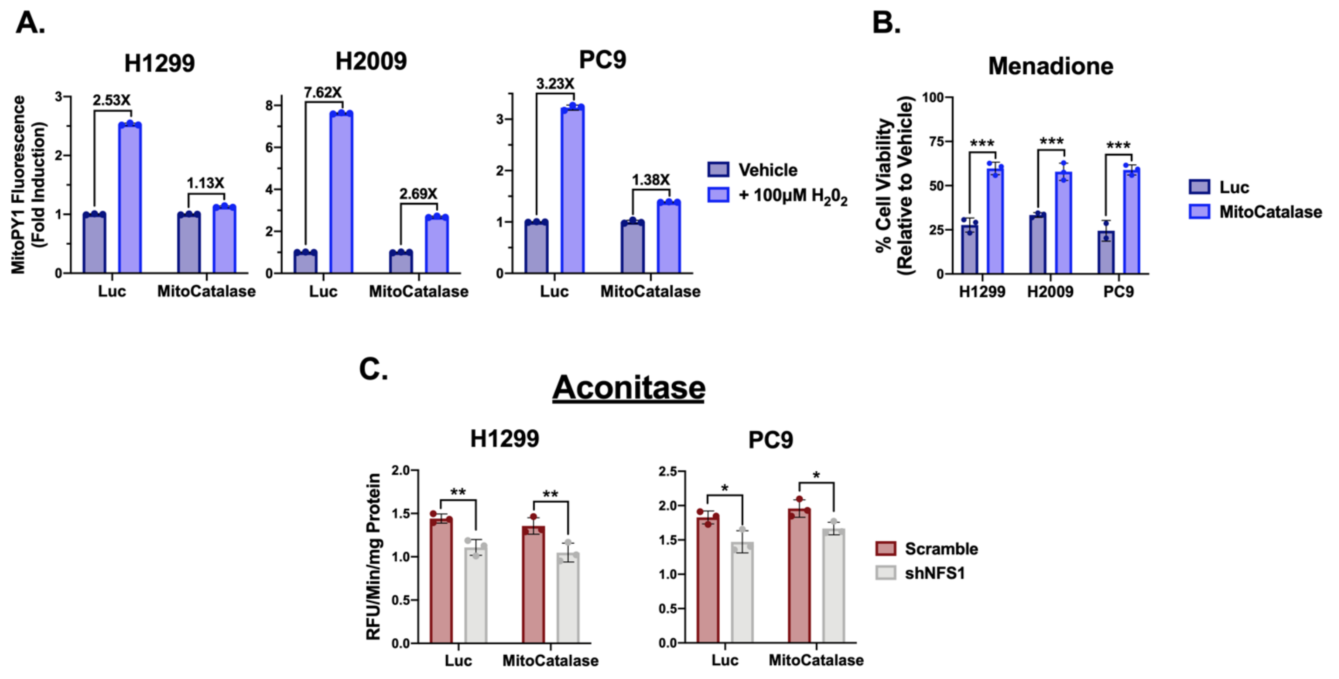


Figure S4. **MitoCatalase protects against mitochondrial oxidation. (A)** Fold inductions of MitoPY1 fluorescence in Luc or MitoCatalase expressing NSCLC cells challenged with 100μM $H_2O_2$. **(B)** Viability of Luc or MitoCatalase expressing NSCLC cells following 24-hour treatment with 15μM (H1299) or 25μM (H2009, PC9) menadione (Student's t test). Cell viability was determined relative to vehicle treated controls. **(C)** Average ACO2 activity in mitochondrial lysates of Luc or MitoCatalase expressing NSCLC cells following NFS1 knockdown (two-way ANOVA). Data are representative of one experiment of three experimental replicates. Data are represented as mean ± SD of three technical replicates. n.s., not significant; *, $p<0.05$; **, $p<0.01$; ***, $p<0.001$.
